# Mindboggling morphometry of human brains

**DOI:** 10.1101/091322

**Authors:** Arno Klein, Satrajit S. Ghosh, Forrest S. Bao, Joachim Giard, Yrjö Häme, Eliezer Stavsky, Noah Lee, Brian Rossa, Martin Reuter, Elias Chaibub Neto, Anisha Keshavan

**Affiliations:** Child Mind Institute, New York, NY, USA; McGovern Institute for Brain Research, Massachusetts Institute of Technology, Cambridge, MA, USA; Department of Otolaryngology, Harvard Medical School, Boston, MA, USA; Department of Electrical and Computer Engineering, University of Akron, Akron, OH, USA; University of Louvain, Belgium; Columbia University, NY, NY, USA; TankThink Labs, Boston, MA, USA; Harvard Medical School, Cambridge, MA, USA; Sage Bionetworks, Seattle, WA, USA; University of California San Francisco, San Francisco, CA, USA

## Abstract

Mindboggle (http://mindboggle.info) is an open source brain morphometry platform that takes in preprocessed T1-weighted MRI data and outputs volume, surface, and tabular data containing label, feature, and shape information for further analysis. In this article, we document the software and demonstrate its use in studies of shape variation in healthy and diseased humans. The number of different shape measures and the size of the populations make this the largest and most detailed shape analysis of human brains every conducted. Brain image morphometry shows great potential for providing much-needed biological markers for diagnosing, tracking, and predicting progression of mental health disorders. Very few software algorithms provide more than measures of volume and cortical thickness, and more subtle shape measures may provide more sensitive and specific biomarkers. Mindboggle computes a variety of (primarily surface-based) shapes: area, volume, thickness, curvature, depth, Laplace-Beltrami spectra, Zernike moments, etc. We evaluate Mindboggle’s algorithms using the largest set of manually labeled, publicly available brain images in the world and compare them against state-of-the-art algorithms where they exist. All data, code, and results of these evaluations are publicly available.

**Author Summary:** Brains vary in many ways, including their shape. Analysing differences in shape between brains or changes in brain shape over time has been used to characterize morphology of diseased brains, but these analyses conventionally rely on simple volumetric shape measures. We believe that access to a greater variety of shape measures could provide greater sensitivity and specificity to morphological disturbances, and could aid in diagnosis, tracking, and prediction of the progression of mental health disorders. Mindboggle is open source software that provides neuroscientists (and indeed, anyone interested in computing shapes) tools for computing a variety of shape measures, including area, volume, thickness, curvature, geodesic depth, travel depth, Laplace-Beltrami spectra, and Zernike moments. In addition to algorithmic contributions, we conducted evaluations and applied Mindboggle to conduct the most detailed shape analysis of human brains.

## 1. Introduction

This article summarizes years of work on the Mindboggle project (http://mindboggle.info), including development and application of software that automates the extraction, identification, and shape analysis of features from human brain magnetic resonance imaging (MRI) data. The principal original contributions of the Mindboggle software include (1) a hybrid approach to combine different software packages’ gray/white matter segmentations, (2) new algorithms for volume and surface shape measures devoted to brain images, including travel depth and cortical thickness, and (3) new shape-based feature extraction algorithms for brain structures such as folds, sulci, and fundi. Further contributions described in this article include (1) evaluations of Mindboggle volume and surface shape measurement algorithms against other software algorithms, (2) evaluation of Mindboggle’s fundus extraction algorithm against other software algorithms, (3) Python implementations of algorithms for general-purpose shape measures such as Laplace-Beltrami spectra and Zernike moments, and (4) application of Mindboggle to provide the most detailed shape measures computed on human brain image data. This Introduction provides background and motivation for the project, Methods outlines the history of the project and the software’s input, processing steps, and output, Results describes evaluations and applications of the software, and Discussion provides commentary and future directions.

### 1.1. The promise of brain imaging for finding biological markers of mental illness

Brain images have been used to derive biological markers of mental illness and disease for years, most notably to predict prognoses among patients with behavioral disorders, often more accurately than current behavioral instruments such as widely used scales and structured interviews. For example, brain images have been used to predict relapse in methamphetamine dependence [1], onset of psychosis in at-risk individuals [2, 3],I recovery from depression eight months later [4], response to drug treatment for depression [5, 6], anxiety [7], and for cognitive behavioral therapy in schizophrenia [8] and social anxiety disorder [9, 10] (see [11] for a more extensive review). Despite the above promising experimental results, there is still a dearth of reliable biomarkers [12]. The importance of identifying new biomarkers is reflected in the National Institute of Mental Health’s Strategic Objectives: “Currently, very few biomarkers have been identified for mental disorders due in part to their complexity and an incomplete understanding of the neurobiological basis of mental disorders…”

### 1.2. Variation in human brains and the “correspondence problem”

A significant impediment to our understanding of mental health is variation in human brain anatomy, physiology, function, connectivity, response to treatment, and so on. The normal range of variation must first be established to determine what is outside of this range, and only then can we hope to address neuropsychiatric assessment, diagnosis, prognosis, treatment, or prevention. An effective biomarker traditionally consists of one or more measures that maximize the separability between groups while minimizing the variance within each group. Brain images provide many ways of measuring different aspects of the brain, but it is not always clear how to compare these measures over time or across individuals. Comparing brains presumes that a brain-to-brain correspondence or mapping has been solved. To do this, scientists ubiquitously co-register images to each other, either individually or in groups, commonly with the use of a standard template brain or labeled atlas. However, registration alone does not guarantee correspondence [13] and templates are often not representative of the group being studied [14, 15]. Additional factors that affect the quality of registration are often ignored. For example, we have empirically demonstrated that registration algorithms vary widely in their accuracy [16], that even the best require removal of non-brain matter to perform adequately [17, 18], and conventional registration is less robust to missing regions than feature-based registration methods [19]. Despite this, many brain imaging studies co-register brains based on image similarity, assume alignment of corresponding anatomy [20], and compare the brains at the level of a small extent such as a sphere or rectilinear volume, which can be on the order of 1/100,000th the volume of the image.

### 1.3. Anatomical feature-based correspondence

Neuroanatomists rely instead on high-level “features” such as distinctive cortical folding patterns and relative positions of subcortical structures to consistently identify anatomical structures or label brain regions ([21, 22]; survey results related to [23]; personal communications with neuroanatomists). Such morphological features may also be identified by using multimodal imaging data and classifiers trained on such data [24]. In addition to whole (gyrus and sulcus) folds, components such as sulcal pits and sulcal fundi hold promise for establishing correspondence across brains. Sulcal pits, points of maximal depth or curvature in sulci, are interesting because they may be well conserved structures formed early in development [25–27] and have been used to characterize conditions such as polymicrogyria [28]. Sulcal fundi are defined as curves that run along the depths of sulci. They form branching skeletons that simplify the complex pattern of folds of the brain, may be measured for morphometry studies, and are used to help define the boundaries between gyri [22]. Like pits, fundi are thought to characterize early stages of morphological development, and therefore may exhibit abnormalities in neurodevelopmental and heritable disorders.

### 1.4. Shape measures as biomarkers

To compare features across individuals we need to quantify them. One quantification method is to characterize the quantities and distributions of grayscale values within a volume, but this does not work well for features of limited extent, such as a point, line, or surface patch. Another method is to coregister a given brain or brain feature with a reference and to define similarity with the reference based on the registration itself (deformation-based morphometry). Yet another method is to directly measure shape, where shape is defined as the geometrical information that remains when location, scale and rotation are removed from an object [29]. Publicly available brain image datasets that include any shape measures usually provide only a few shape measures per anatomical region: volume (such as the Internet Brain Volume Database, http://www.nitrc.org/projects/ibvd), surface area, and/or cortical thickness. These measures are useful for studies of neurogenesis or atrophy in morphological development, degeneration, injury, and disease progression. Volume measurement is almost ubiquitous in such studies, and cortical thickness measures derived from structural MRI data have been reported to help characterize a variety of disorders [30] such as mild cognitive impairment and Alzheimer’s disease [31–33], multiple sclerosis [34], schizophrenia [35], autism spectrum disorder [36], and alcohol dependence [37], and to predict onset or progression of, for example, Alzheimer’s disease [38–44], major depressive disorder [45], and attention-deficit/hyperactivity disorder [46].

More subtle shape measures may provide more sensitive and specific biomarkers, and combining shape measures in a multivariate analysis can improve results over any single measure [47]. The lack of shape measures may be attributable to the paucity of software programs such as BrainVisa [48, 49] (https://www.nitrc.org/projects/brainvisa_ext) that compute more nuanced measures. Sulcal width has been used to differentiate between groups with mild cognitive impairment [50] and global and local gyrification indices computed from sulci have been used to characterize schizophrenia [51] and early-onset vs. intermediate-onset bipolar disorder as well as bipolar and unipolar depression [52–54]. More abstract shape measures such as Zernike moments (see below) have been used in patient classification, such as to distinguish cases of dementia [55].

## 2 Methods

Mindboggle (http://mindboggle.info) is an open source brain morphometry platform that takes in preprocessed T1-weighted MRI data, and outputs volume, surface, and tabular data containing label, feature, and shape information for further analysis. Mindboggle can be run on the command line as “mindboggle” and also exists as a cross-platform Docker container for convenience and reproducibility of results. The software runs on Linux and is written in Python 3 and Python-wrapped C++ code called within a modular Nipype pipeline framework (http://nipy.org/nipype,doi:10.5281/zenodo.50186) to promote a modular, flexible design that captures provenance information [56]. We have tested the software most extensively with Python 3.5.1 on Ubuntu Linux 14.04. Issues and bugs are tracked on GitHub (https://github.com/nipy/mindboggle/issues) and support questions are posted on NeuroStars (https://neurostars.org/t/mindboggle/) with the tag “*mindboggle*”.

Mindboggle’s flexible, modular, open source pipeline facilitates the addition of functions for computing almost any shape measure in any programming language. We initialized Mindboggle with shape measures that we thought have great potential for describing the shapes of brain structures and that complement shape measures supplied by existing software packages. It is just as easy to include functions in Mindboggle for volume-based as it is for surface-based measures, but we decided to focus primarily on surface-based shape measures to complement the volume-based methods available in standard brain image analysis packages. We also want to emphasize in this work intrinsic shape measures of brain structures rather than shapes inferred by registration-based methods such as voxel-based, tensor-based, and deformation-based morphometry that rely on a reference or canonical template and are sensitive to errors in registration. We also do not consider density values to be intrinsic shape measures, as they do not describe the shape of an object, but quantify values obtained within an object, in an analogous manner as one would quantify an fMRI signal or PET ligand binding within a voxel or region of interest.

### 2.1 History of the Mindboggle open source brain morphometry platform

**2005:** The initial version of the Mindboggle software (https://osf.io/gfwcn/) was written in Matlab (Mathworks, Inc., Natick, MA) as part of a doctoral dissertation [57]. It introduced a feature-driven approach to label human brain MRI data using one atlas [19] or multiple atlases [58].

**2009:** With generous funding from the National Institute of Mental Health, we began to write Mindboggle from scratch in Python with some surface mesh measurements programmed in C++, all run from a software pipeline written in the Nipype framework.

**2010:** To ensure that the most consistent and accurate anatomical labels are assigned to brain image data, we introduced a new cortical labeling protocol with 62 labels (Fig 1) called the Desikan-Killiany-Tourville (DKT) protocol [22, 23] (http://mindboggle.info/labels.html). We applied this protocol to manually edit the anatomical labels for 101 individuals (20 of which also include CMA non-cortical labels [59] (http://www.cma.mgh.harvard.edu/manuals/segmentation/)). The resulting Mindboggle-101 dataset [22, 60] (http://mindboggle.info/data.html, http://mindboggle.info/data.html, https://osf.io/nhtur/) is still the largest publicly available set of manually edited human brain labels in the world. These brains were used to construct multiple templates [61] and atlases [62], including the joint fusion [63] volume atlas (https://osf.io/d2cmy/?action=download&version=1) used by the Mindboggle software for volume-based segmentation and labeling, and the DKT-40 and DKT-100 surface atlases [62] used for labeling cortical surfaces by the FreeSurfer software package [64–66] (https://surfer.nmr.mgh.harvard.edu/fswiki). The DKT-100 is used as the default atlas for labeling brains in FreeSurfer (version 6). The Mindboggle-101 brains are used for evaluations and shape analyses described in the Results section.

**2013:** A prototype for online, interactive visualization of Mindboggle shape analysis data won a hackathon challenge at the Human Brain Mapping (HBM 2013) conference. After use of the XTK (https://github.com/xtk/X#readme) WebGL JavaScript library [67, 68], we used the threejs (http://threejs.org/) and D3 JavaScript libraries in a second (HBM 2015 [69]) and third (HBM 2016) hackathon to create the ROYGBIV online interactive brain image viewer (Fig 1; http://roygbiv.mindboggle.info), which is under active development (https://github.com/binarybottle/roygbiv).

**Fig 1.**
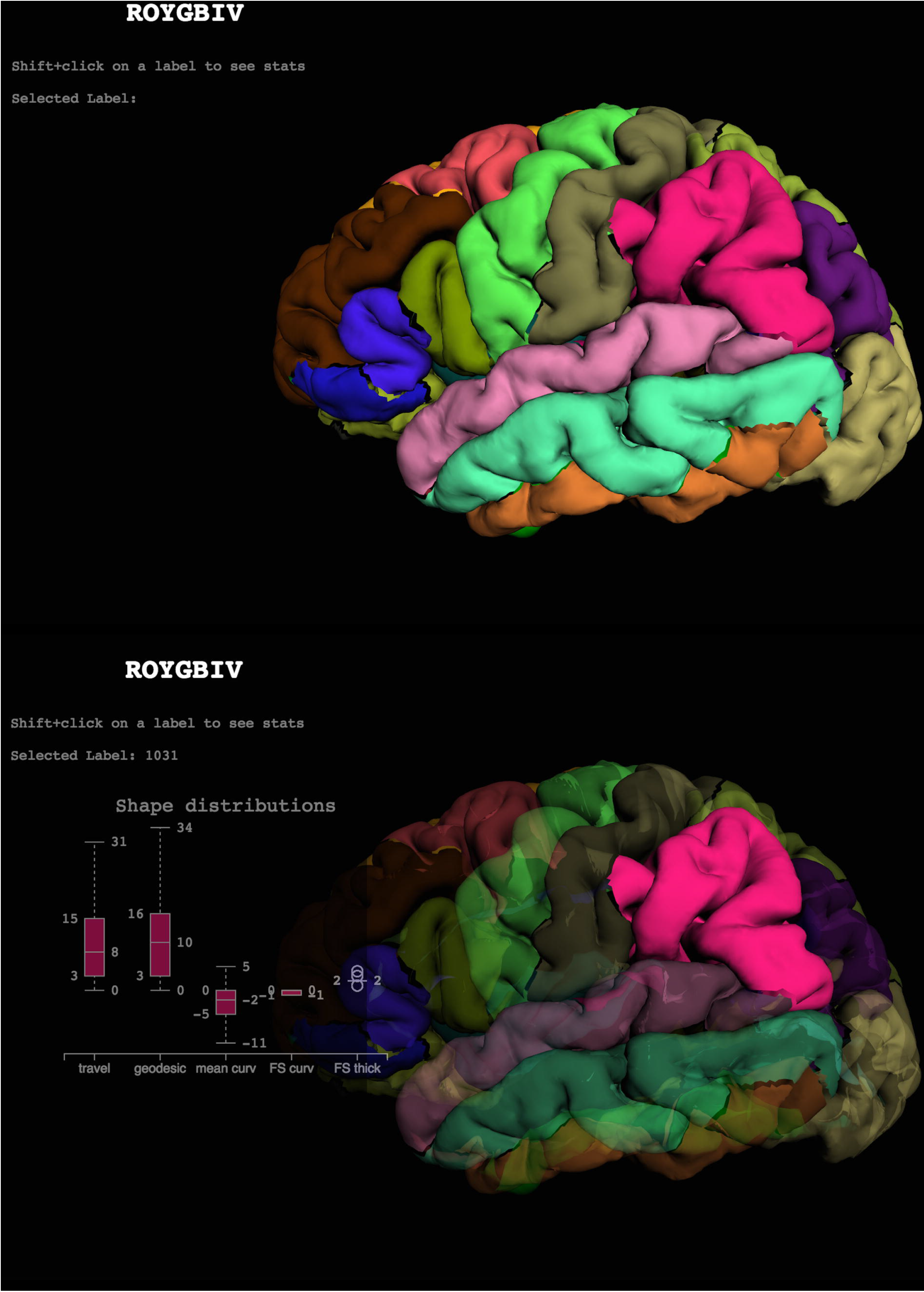
Cortical labels displayed in the ROYGBIV interactive online brain image viewer. The anatomical labels included in the DKT cortical labeling protocol [22] used to label the Mindboggle-101 data are displayed on a left cortical surface. These two panels show the current state of our prototype for a browser-based interactive visualization of the left hemisphere of a human brain [69] and accompanying plot of some of Mindboggle’s shape measures for a selected region (http://roygbiv.mindboggle.info).

**2015:** Mindboggle processed Alzheimer’s Disease Neuroimaging Initiative (ADNI; adni.loni.usc.edu; [70]) and AddNeuroMed [71] data for an international Alzheimer’s disease challenge [72] (https://www.synapse.org/Synapse:syn2290704/wiki/60828). Teams performed statistical analyses on Mindboggle shape measures to try and determine which brains had Alzheimer’s disease, mild cognitive impairment, or were healthy, and to try and estimate a cognitive measure (mini-mental state exam score). The Results section presents an analysis of some of these data.

**2016:** Mindboggle is launched for broader public use after making the following improvements:

- Software ported from Python 2 to Python 3
- Docstring tests provided for almost every function
- GitHub repository transferred to the nipy.org community’s GitHub account
- Online documentation updated automatically
- Online support via NeuroStars with the tag *“mindboggle”*
- Online tests run automatically

The documentation is updated online (https://readthedocs.org/projects/mindboggle) and the tests are updated online (https://circleci.com/gh/nipy/mindboggle) every time a commit is made to the GitHub repository (https://github.com/nipy/mindboggle).

#### 2.3 Input data and preprocessing

For running individual functions on surface meshes, the only inputs to the software are outer cortical surface meshes constructed from T1-weighted MRI data by software such as FreeSurfer, Caret [73] or BrainVISA [48], once converted to an appropriate format (see below). For this study we used FreeSurfer v5.1-derived labels and meshes, but the recently released FreeSurfer version 6 is recommended because it uses Mindboggle’s DKT-100 surface-based atlas (with the DKT31 labeling protocol) by default to generate labels on the cortical surfaces, and generates corresponding labeled cortical and non-cortical volumes (wmparc.mgz) [74]. To preprocess data for use by Mindboggle, run the following FreeSurfer command on a T1-weighted `$IMAGE` file (e.g., subject1.nii.gz) to output a `$SUBJECT` folder (e.g., subject1):

`recon-all -all -i $IMAGE -s $SUBJECT`

The `recon-all` command performs many steps (https://surfer.nmr.mgh.harvard.edu/fswiki/recon-all), but the ones that are most relevant include (1) segmentation of the brain image into different tissue classes (gray/white/cerebrospinal fluid), (2) reconstruction of a triangular surface mesh approximating the pial surface for each brain hemisphere, and (3) anatomical labeling of each surface and each volume.

To refine segmentation, labeling, and volume shape analysis, Mindboggle optionally takes output from the Advanced Normalization Tools (ANTs, v2.1.0rc3 or higher recommended; http://stnava.github.io/ANTs/), which performs various image processing steps such as brain volume extraction [17, 75], tissue-class segmentation [76], and registration-based labeling [16,18,75]. To generate the ANTs transforms and segmentation files used by Mindboggle, run the `antsCorticalThickness.sh` script [75] on the same `$IMAGE` file, set an output `$PREFIX`, and provide paths to the OASIS-30 Atropos template files in directory `$TEMPLATE` (backslash denotes a line return):

~~~
antsCorticalThickness.sh -d 3 -a IMAGE -o PREFIX \
     -e $TEMPLATE/T_template0.nii.gz \
     -t $TEMPLATE/T_template0_BrainCerebellum.nii.gz \
     -m $TEMPLATE/T_template0_BrainCerebellumProbabilityMask.nii.gz \
     -f $TEMPLATE/T_template0_BrainCerebellumExtractionMask.nii.gz \
     -p $TEMPLATE/Priors2/priors%d.nii.gz
~~~

Links to the template (https://osf.io/bx35m/?action=download&version=1) and example input data (https://osf.io/8cf5z/) can be found on the Mindboggle website.

#### 2.3 Mindboggle processing steps

The following steps are performed by Mindboggle (http://mindboggle.info/software.html):

1. Convert FreeSurfer formats to NIfTI volumes and VTK surfaces.
2. Optionally combine FreeSurfer and ANTs gray/white segmented volumes and fill with labels.
3. Compute volumetric shape measures for each labeled region.
4. Compute shape measures for every cortical surface mesh vertex.
5. Extract cortical surface features.
6. Segment cortical surface features with labels.
7. Compute shape measures for each cortical surface label or sulcus.
8. Compute statistics for each shape measure in Step 4 for collections of vertices.

##### Step 1: Convert FreeSurfer formats to NIfTI volumes and VTK surfaces

Mindboggle performs all of its processing in two open standard formats: NIfTI (.nii.gz; http://nifti.nimh.nih.gov/) for volume images and VTK (.vtk, Visualization Toolkit; http://www.vtk.org/) for surface meshes. ANTs output already supports NIfTI; given FreeSurfer input, the first step that Mindboggle performs is to convert FreeSurfer volume and surface formats to NIfTI and VTK for further processing. All volume images in this study have a resolution of 1x1x1 mm^3^ per voxel (volume element). All surface-based shape measures are computed on the “pial surface” (cortical-cerebrospinal fluid boundary) by default, since it is sensitive to differences in cortical thickness.

##### Step 2: Optionally combine FreeSurfer and ANTs gray/white segmented volumes and fill with labels

This optional step of the pipeline will be skipped in the future when methods for tissue class segmentation of T1-weighted MR brain images into gray and white matter improve. FreeSurfer and ANTs make different kinds of mistakes while performing tissue class segmentation (Fig 2). After visual inspection of the gray/white matter boundaries in over 100 EMBARC (http://embarc.utsouthwestern.edu/, http://embarc.utsouthwestern.edu/, https://clinicaltrials.gov/ct2/show/NCT01407094) brain images processed by FreeSurfer, we found that at least 25 brains had significant overcropping of the brain, particularly in ventral regions such as lateral and medial orbitofrontal cortex and inferior temporal lobe due to poor surface mesh reconstruction in those regions. This corroborates Klauschen’s observation that FreeSurfer underestimates gray matter and overestimates white matter [77]. We also found that ANTs tends to include more cortical gray matter than FreeSurfer, but at the expense of losing white matter that extends deep into gyral folds, and sometimes includes non-brain tissue such as transverse sinus, sigmoid sinus, superior sagittal sinus, and bony orbit.

Mindboggle attempts to reconcile the differences between FreeSurfer and ANTs segmentations by combining them. The `relabel_volume` function converts the (wmparc.mgz) labeled file generated by FreeSurfer and the (BrainSegmentation.nii.gz) segmented file generated by the ANTs `Atropos` function [76] to binary files of pseudo-white matter and gray (including deep gray) matter. The `combine_2labels_in_2volumes` function overlays FreeSurfer white matter atop ANTs cortical gray, by taking the union of cortex voxels from both binary files as gray matter, the union of the non-cortex voxels from the two binary files as white matter, and assigning intersecting cortex and non-cortex voxels as non-cortex. While this strategy often preserves gray matter bordering the outside of the brain, it still suffers from over-inclusion of non-brain matter, and sometimes replaces true gray matter with white matter in areas where surface reconstruction makes mistakes.

The FreeSurfer/ANTs hybrid segmentation introduces new gray-white matter boundaries, so the corresponding anatomical (gyral-sulcal) boundaries generated by FreeSurfer and ANTs need to be updated accordingly. Mindboggle uses `ImageMath’s PropagateLabelsThroughMask` function in ANTs to propagate both FreeSurfer and ANTs anatomical labels to fill the gray and white matter volumes independently. The FreeSurfer-labeled cerebellum voxels overwrite any intersecting cortex voxels, in case of overlap.

**Fig 2.**
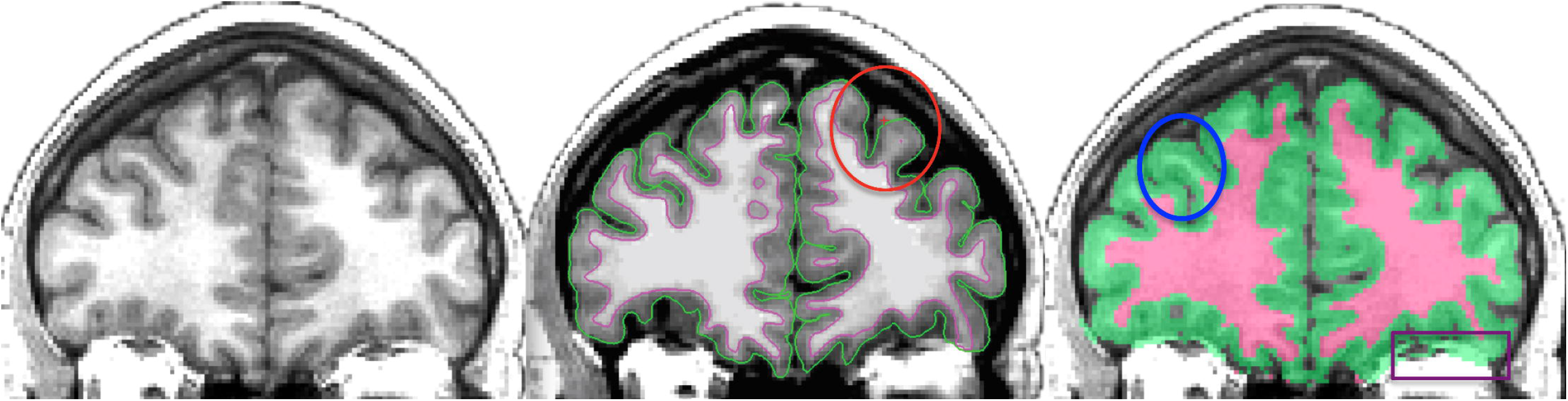
FreeSurfer and ANTs gray/white matter segmentation. Left: Coronal slice of a T1-weighted brain MRI. Middle: Cross-section of FreeSurfer inner (magenta) and outer (green) cortical surfaces overlaid on top of the same slice. The red ellipse circumscribes a region where the FreeSurfer surface reconstruction failed to include gray matter on the periphery of the brain. Right: Cross-section of ANTs segmentation. The blue ellipse circumscribes a region where the ANTs segmentation failed to segment white matter within a gyrus that the FreeSurfer correctly segmented (compare with the middle panel). The purple box in the lower right highlights a region outside of the brain that the ANTs segmentation mistakenly includes as gray matter. To reconcile some of these discrepancies, Mindboggle currently includes an optional processing step that combines the segmentations from FreeSurfer and ANTs. This step essentially overlays the white matter volume enclosed by the magenta surface in the middle panel atop the gray/white segmented volume in the right panel.

##### Step 3: Compute volumetric shape measures for each labeled region

- volume
- thickness of cortical labels `(thickinthehead)`

As mentioned in the Introduction, the most common shape measures computed for brain image data are volume and cortical thickness for a given labeled region of the brain. Volume measurements are influenced by various factors such as cortical thickness, surface area [78], and microstructural tissue properties [79]. Computing the volume per labeled region is straightforward: Mindboggle’s `volume_per_brain_region` function simply multiplies the volume per voxel by the number of voxels per region. In contrast, cortical thickness can be estimated using a variety of MRI processing algorithms [49,75,80–83]. Since Mindboggle accepts FreeSurfer data as input, we include FreeSurfer cortical thickness [80] estimates with Mindboggle’s shape measures. When surface reconstruction from MRI data produces favorable results (see above), FreeSurfer cortical thickness measures can be highly reliable [81,84,85]. See Results for our evaluation of cortical thickness measures.

To avoid surface reconstruction-based problems with the cortical thickness measure, we built a function called `thickinthehead` that computes a simple thickness measure for each cortical region from a brain image volume without relying on surface data (Fig 3). The `thickinthehead` function first saves a brain volume that has been segmented into cortex and non-cortex voxels into separate binary files, then resamples these cortex and non-cortex files from, for example, 1mm^3^ to 0.5mm^3^ voxel dimensions to better represent the contours of the cortex. Next it extracts outer and inner boundary voxels of the cortex by morphologically eroding the cortex by one (resampled) voxel bordering the outside of the brain and bordering the inside of the brain (non-cortex). Then it estimates the middle cortical surface area by the average volume of the outer and inner boundary voxels of the cortex.

Finally, it estimates the thickness of a labeled cortical region as the volume of the labeled region divided by the middle surface area of that region. The `thickinthehead` function calls the `ImageMath`, `Threshold`, and `ResampleImageBySpacing` functions in ANTs.

**Fig 3.**
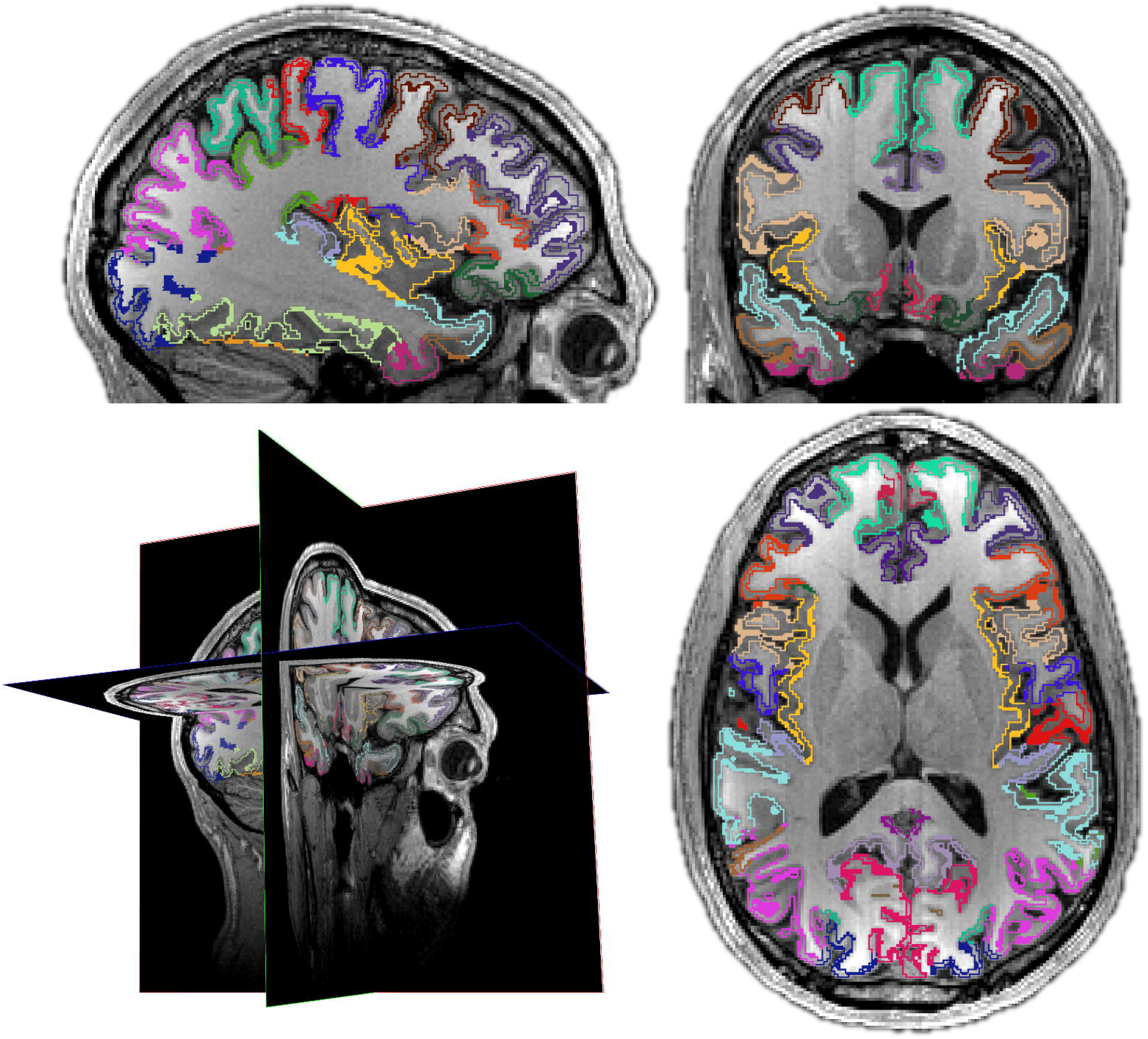
Thickinthehead estimates average cortical thickness per brain region. Mindboggle’s `thickinthehead` algorithm estimates cortical thickness for each brain region without relying on cortical surface meshes by dividing the volume of a region by an estimate of its middle surface area. Clockwise from lower left: 3-D cross-section and sagittal, coronal, and axial slices. The colors represent the inner and outer “surfaces” of cortex created by eroding gray matter bordering white matter and eroding gray matter bordering the outside of the brain. The middle surface area is estimated by taking the average volume of these inner and outer surfaces.

##### Step 4: Compute shape measures for every cortical surface mesh vertex

- surface area
- mean curvature
- geodesic depth
- travel depth
- convexity (FreeSurfer)
- thickness (FreeSurfer)

Aside from the convexity and thickness measures inherited from FreeSurfer, shape measures computed for each vertex of a cortical surface triangular mesh are generated by Mindboggle’s open source C++ code (using the Visualization Toolkit, VTK) developed by Joachim Giard: surface area, mean curvature, geodesic depth, and travel depth. Surface area is computed per vertex (as opposed to per face of the mesh to be consistent with all other Mindboggle shape measures) as the area of the Voronoi polygon enclosing the vertex (Fig 4). Area can be used to normalize other values computed within a given region such as a gyrus or sulcus [86].

**Fig 4.**
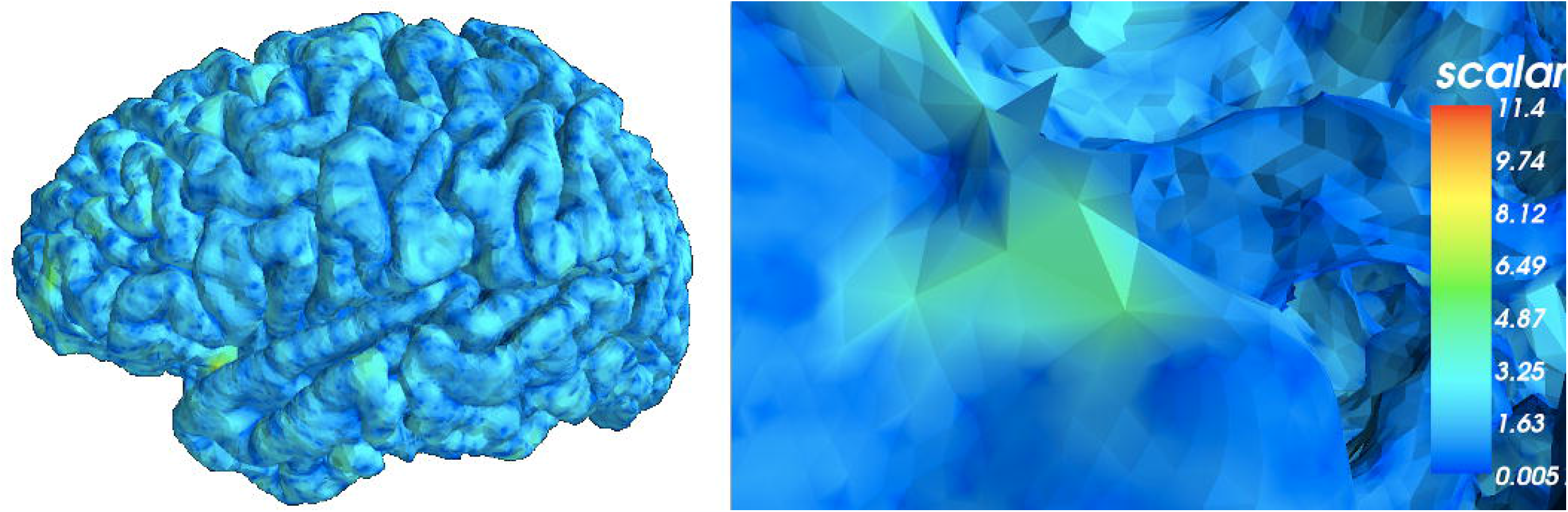
Surface area per vertex. Mindboggle computes surface area for each surface mesh vertex as the area of the Voronoi polygon enclosing the vertex. Left: Lateral view of a left cortical hemisphere colored by surface area per vertex. Right: Closeup of the surface mesh. Mindboggle uses area to normalize other shape values computed within a given region such as a gyrus or sulcus.

Curvature is an obvious shape measure for a curved and folded surface like the cerebral cortex and has the potential to help make inferences about other characteristics of the brain, such as sulcus width, atrophy [87, 88], structural connections [89] and differential expansion of the cortex [90]. Mindboggle computes both mean and Gaussian curvatures based on the relative direction of the normal vectors in a small neighborhood (Fig 5), which works best for low resolution or for local peaks, but can be sensitive to the local linear geometry of the mesh. Increasing the radius of the neighborhood mitigates this sensitivity, so a neighborhood parameter corresponding to the radius of a geodesic disk is defined in the unit of the mesh. If coordinates are in millimeters, the default setting of 2 results in an analysis of the normal vectors within a 2mm radius disk. Other options include computing both mean and Gaussian curvatures based on the local ratios between a filtered surface and the original surface area (the filtering is done using Euclidean distances, so it’s best for less accurate but fast visualization), or computing the mean curvature based on the direction of the displacement vectors during a Laplacian filtering (a good approximation based on the Laplacian, but underestimates very large, negative or positive, curvatures due to saturation).

**Fig 5.**
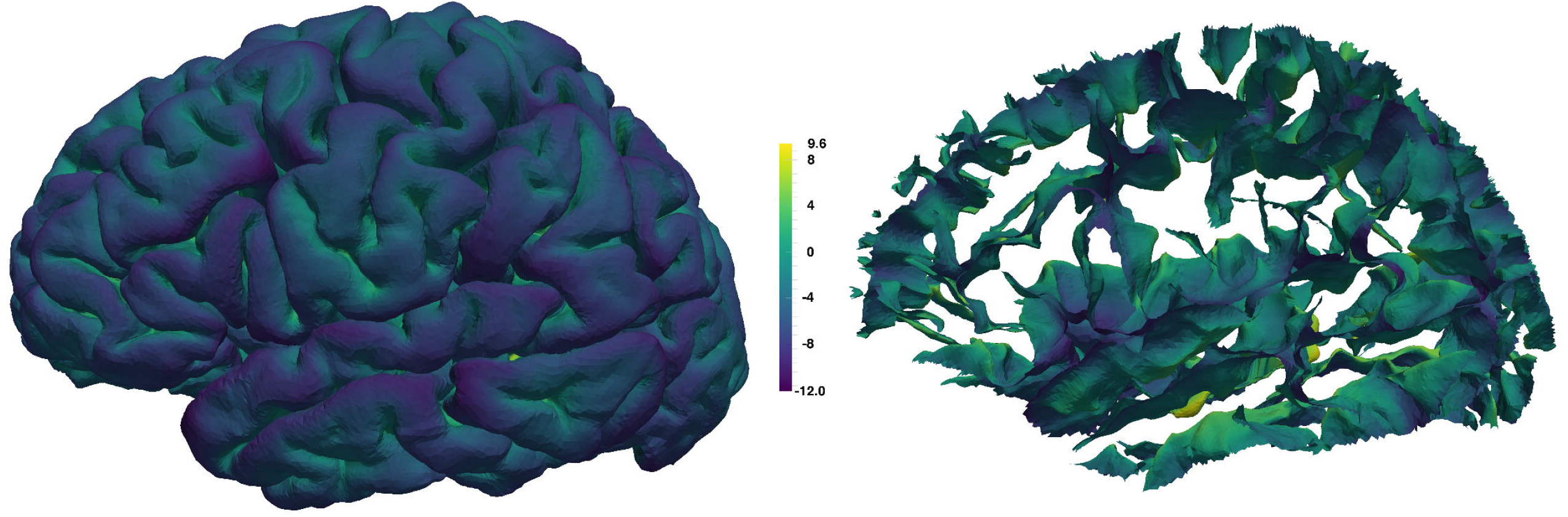
Curvature per vertex. Mindboggle computes curvature for each surface mesh vertex. This medial view of the left cortical hemisphere is colored by curvature per vertex (green indicates flat regions).

Depth is an important measure characterizing the highly folded surface of the human cerebral cortex. Since much of the surface is buried deep within these folds, an accurate measure of depth is useful for defining and extracting deep features, such as sulci [91, 92], sulcal fundus curves [93–95], and sulcal pits [26,96,97]. Depth may also serve as an indicator of developmental stage [26].

We are aware of three predominant methods for measuring depth of points on the surface of the cerebral cortex, where depth is the distance between a given point on the brain surface to an outer reference surface of zero depth (the portions of the brain surface in contact with the outer reference surface are gyral crowns or crests). The first is Euclidean depth, the distance along a straight path from the point on the brain to the outer reference surface. A straight path has the undesirable property that it will cross through anything, which can make a highly folded surface indistinguishable from a slightly folded surface that fills the same volume. The second is geodesic depth, the shortest distance along the surface of the brain from the point to where the brain surface makes contact with the outer reference surface.

Geodesic paths are very sensitive to slight or gradual changes in depth, resulting in exaggerated distances where the outer reference surface does not wrap the brain closely. Geodesic paths are also greatly affected by cavities, so distances can be exaggerated where there are irregularities, particularly in the bottoms of sulcus folds. The third measure, FreeSurfer software’s “convexity,” while not explicitly referred to as depth, is used to indicate relative depth. It is based on the displacement of surface mesh vertices after inflating the surface mesh [64]. This can result in assigning positive depth to points on the outermost surface of the brain such as on a gyral crest, however, which is not consistent with an intuitive measure of depth.

Travel depth was introduced as a hybrid depth measure for macromolecules, defined as the shortest distance that a solvent molecule would travel from the convex hull of the macromolecule without penetrating the macromolecule surface. It was first defined for surfaces but using a voxel-based algorithm [98] that uses Dijkstra’s algorithm for finding shortest paths, and was later refined to use a much faster and more accurate vertex-based computation [99]. A detailed description of the latter follows. The travel depth algorithm constructs a combination of Euclidean paths outside the cortical surface and estimated geodesic paths along the cortical surface. The principal idea of the algorithm lies in the classification of a surface into “visible” and “hidden” areas (Fig 1 in **Supplement 1**). A point on the surface is considered “visible” by another point if they can be connected by a straight line without intersecting the volume enclosed by the surface. In other words, there is a “line of sight” between the two points that does not run through the interior of the surface. A point is considered “hidden” from another point if it is not visible and can only be reached by a path running either along the surface or connecting points of the surface without intersecting the enclosed volume.

The above implementations of travel depth use a convex hull (Fig 2 in **Supplement 1**), as do most measures of cortical depth such as the adaptive distance transform [100], while other algorithms do not define a zero-depth reference surface but rely instead on convergence of an algorithm, such as the depth potential map [101]. The shape of the brain is concave in places, resulting in some gyral crowns that do not touch the convex hull. For example, in Fig 3 in **Supplement 1**, the gyri of the medial temporal lobe are assigned positive depth, resulting in an unreasonably high depth for the folds of that region. Since the convex hull is not suitable for application to brain images, or for surfaces with global concavities, we define and construct a different reference surface that we call the wrapper surface (Fig 5 in **Supplement 1**). The wrapper surface has to be chosen such that the top of a gyrus has zero depth. We compute a wrapper surface as follows. We create a volume image representing the interior of the mesh, dilate this image with a probe of radius *r*, then erode it with the same probe. This operation is also known as morphological closing, and it is important to carefully set the probe radius. If the radius is too large, the wrapper surface will be similar to the convex hull, and if the radius is too small, the wrapper surface will be too close to the original surface and the travel depth will be close to zero even inside folds. We used an empirically determined radius of 5 mm. The wrapper surface mesh is an isosurface of this morphologically closed image volume, created using the marching cubes algorithm. On a brain mesh with 150,000 vertices and 300,000 triangles, the algorithm takes around 200 seconds on an ordinary computer when the wrapper surface is provided. The generation of the wrapper surface takes an additional 20 seconds for a probe radius of 5 mm.

Mindboggle’s travel depth algorithm assigns a depth value to every vertex in a mesh, is faster and more accurate than voxel-based approaches, assigns more reasonable path distances that are less sensitive to surface irregularities and imaging artifacts than geodesic distances, and is faithful to the topology of the surface. Fig 6 shows an example of geodesic and travel depth values, and the Results section summarizes our comparison of travel depth with geodesic depth and FreeSurfer convexity measures.

**Fig 6.**
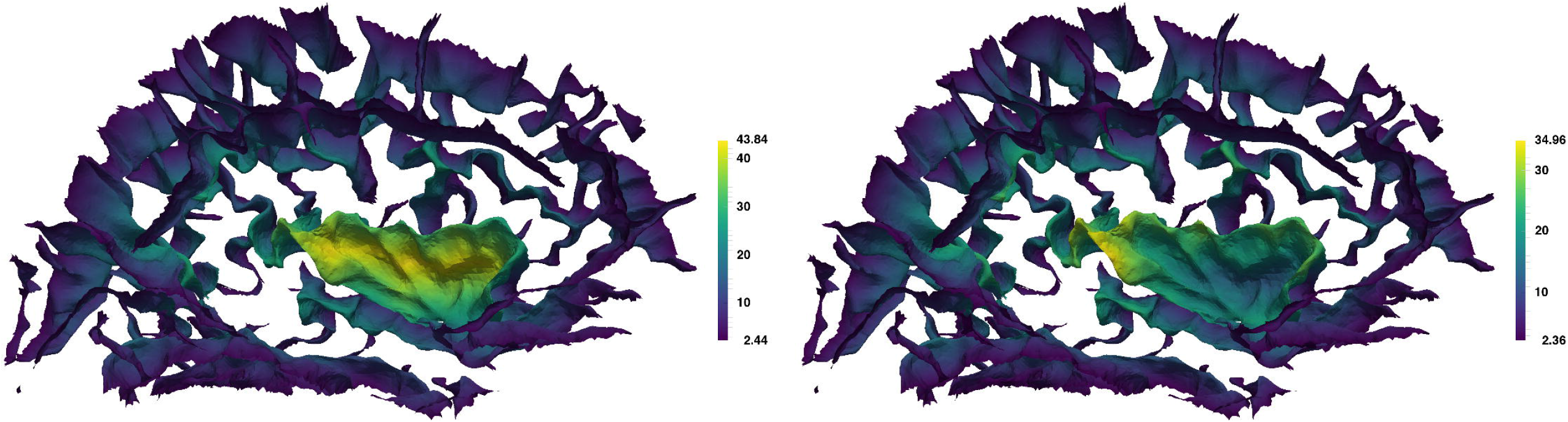
Geodesic depth and travel depth per vertex. Mindboggle computes geodesic depth (left) and travel depth (right) for each surface mesh vertex. This medial view of the sulcus folds from the left cortical hemisphere is colored by depth, with the deepest vertices in yellow. Note that the deepest vertices according to geodesic depth reside toward the center of the insula (center fold), whereas the deepest vertices according travel depth run along the deepest furrows of the insula, as one would expect.

##### Step 5: Extract cortical surface features

- folds
- fundus per fold

Mindboggle extracts hierarchical structures from cortical surfaces [102, 103], including folds and fundus curves running along the depths of the folds. A fold is a group of connected, deep vertices (left side of Fig 7). When assigned anatomical labels, folds can be broken up into sulci (right side of Fig 7). To extract folds, a depth threshold is used to segment deep vertices of the surface mesh. We have observed in the histograms of travel depth measures of cortical surfaces that there is a rapidly decreasing distribution of low depth values (corresponding to the outer surface, or gyral crowns) with a long tail of higher depth values (corresponding to the folds). Mindboggle’s `find_depth_threshold` function therefore computes a histogram of travel depth measures, smooths the histogram’s bin values, convolves to compute slopes, and finds the depth value for the first bin with zero slope. The `extract_folds` function uses this depth value, segments deep vertices, and removes extremely small folds (empirically set at 50 vertices or fewer out of a total mesh size of over 100,000 vertices).

**Fig 7.**
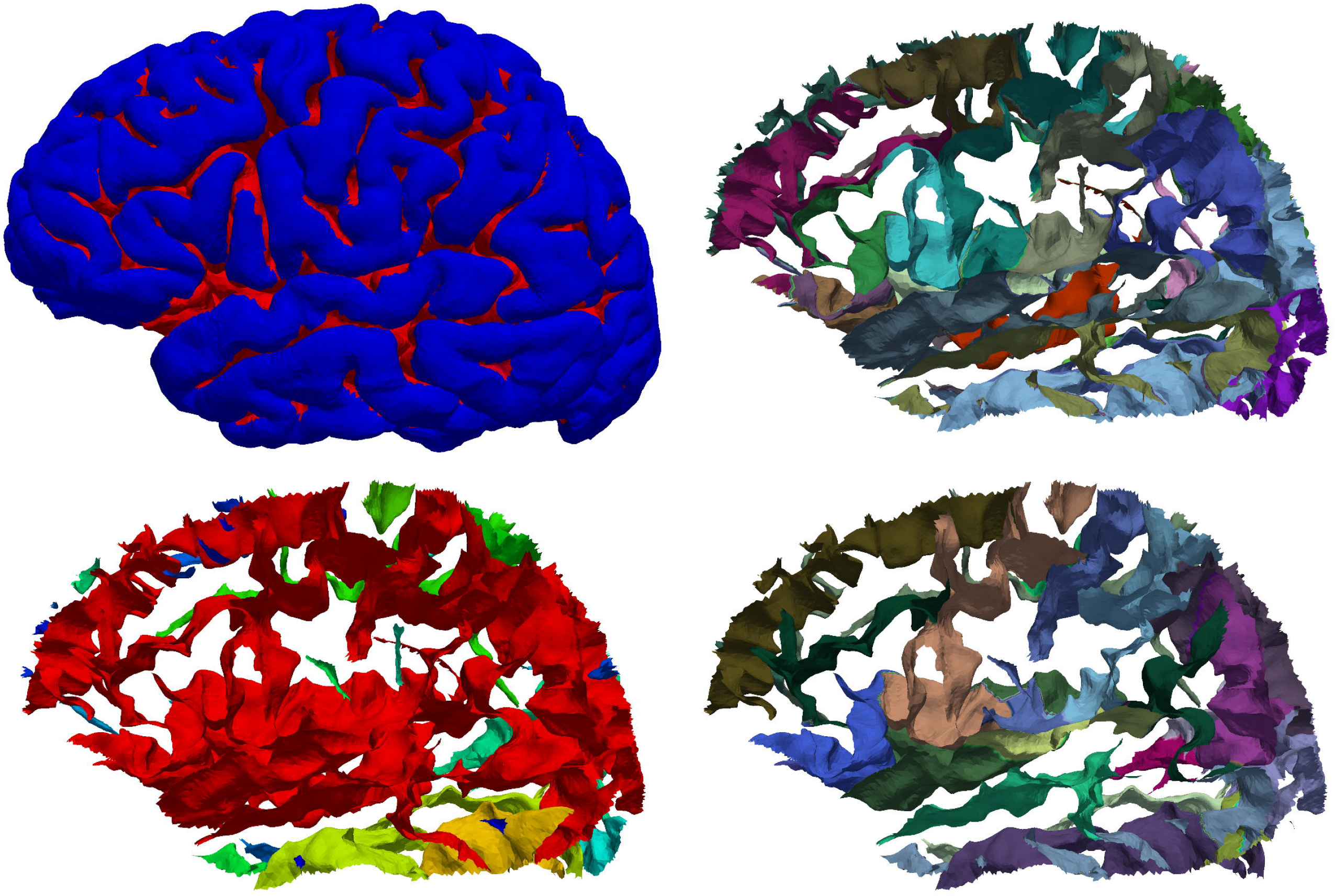
Cortical fold extraction and sulcus segmentation. Top left: Lateral view of the left hemisphere of a brain with folds labeled red. Mindboggle extracts cortical surface folds based on a depth threshold that it computes from the distribution of travel depth values. Bottom left: individually colored folds from the same brain. The red surface shows that folds can be broadly connected, depending on the depth threshold, and therefore do not map one-to-one to anatomical region labels. Top right: The same folds with individually colored anatomical labels. These labels can be automatically or manually assigned (as in the case of this Mindboggle-101 subject). Bottom right: Individually colored sulci. Mindboggle uses the anatomical labels to segment folds into sulci, defined as folded portions of cortex whose opposing banks are labeled with sulcus label pairs in the DKT labeling protocol [22]. Each label pair is unique to one sulcus and represents a boundary between two adjacent gyri, so sulcus labels are useful to establish correspondences across brains. Portions of folds that are missing are not defined as sulci by the DKT labeling protocol.

A fundus is a branching curve that runs along the deepest and most highly curved portions of a fold (Fig 8). As mentioned above, fundi can serve as boundaries between anatomical regions and are interesting for their relationship to morphological development and disorders. But they are too tedious, time-consuming, and difficult to be drawn in a consistent manner on the surface meshes derived from MR images. Mindboggle provides multiple functions for extracting fundi which are optionally generated from the command line; the `extract_fundi` function described below is used by default and is evaluated against other fundus extraction methods in the Results section. This function extracts one fundus from each fold by finding the deepest vertices inside the fold, finding endpoints along the edge of the fold, connecting the former to the latter with tracks that run along deep and curved paths (through vertices with high values of travel depth multiplied by curvature), and running a final filtration step. A more detailed description of these four steps follows. In the first step, the deepest vertices are those with values at least two median absolute deviations above the median (non-zero) value. If two of these deep vertices are within (a default of) 10 edges from each other, the vertex with the higher value is chosen to reduce the number of possible fundus paths as well as to reduce computation time. To find the endpoints in the second step, the `find_outer_endpoints` function propagates multiple tracks from seed vertices at median depth in the fold through concentric rings toward the fold’s edge, selecting maximal values within each ring, and terminating at candidate endpoints. The final endpoints are those candidates at the end of tracks that have a high median value. If two candidate endpoints are within (a default of) 10 edges from each other, the endpoint with the higher value is chosen; otherwise the resulting fundi can have spurious branching at the fold’s edge. The `connect_points_erosion` function connects the deepest fold vertices to the endpoints with a skeleton of 1-vertex-thick curves by erosion. It erodes by iteratively removing simple topological points and endpoints in order of lowest to highest values, where a simple topological point is a vertex that when added to or removed from an object on a surface mesh (such as a fundus curve) does not alter the object’s topology.

**Fig 8.**
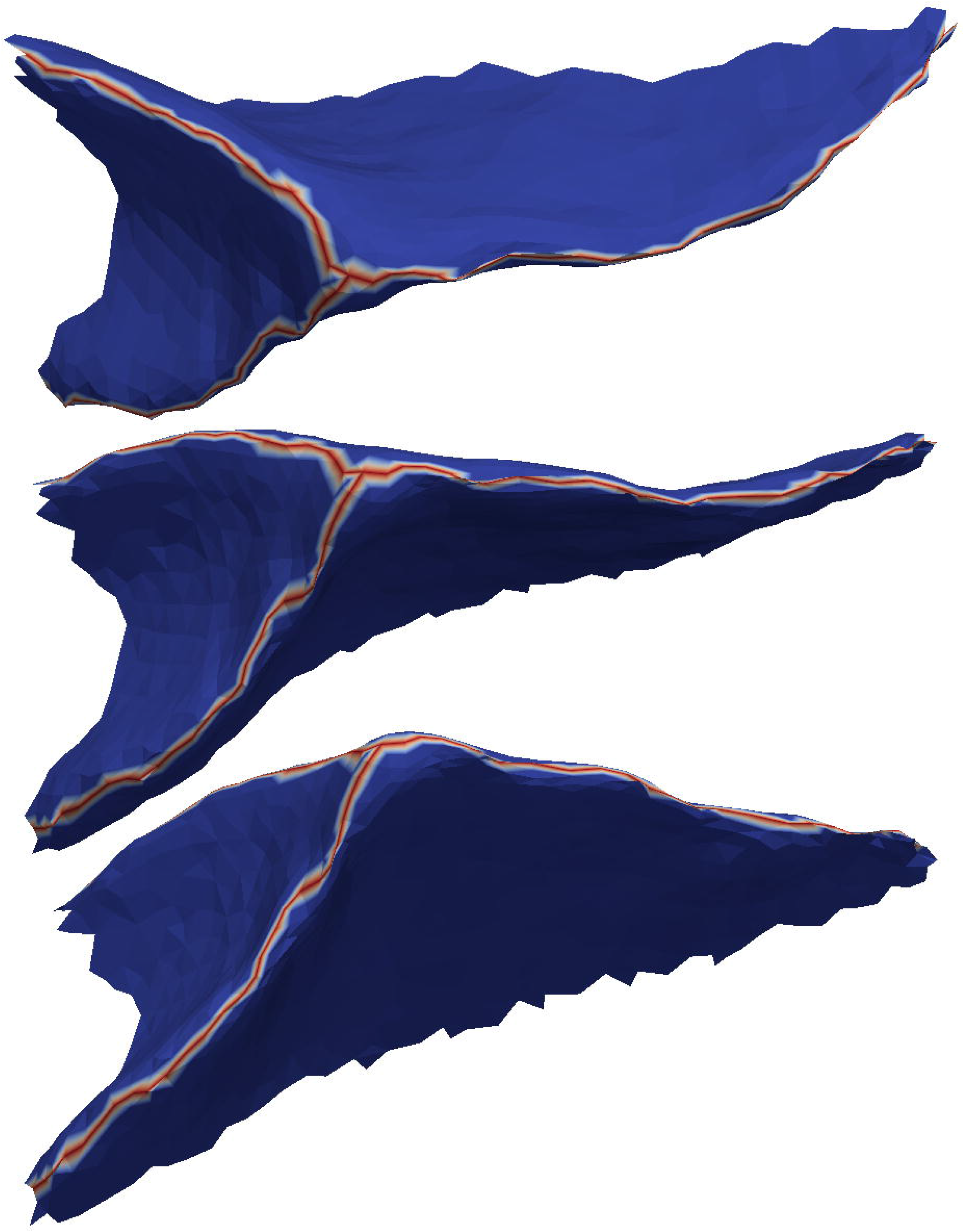
Sulcal fundi. This figure shows three views of the outside of a single sulcus (taken from the top middle fold in Fig 7) to clearly show a simple example of a fundus (red branching curve). Mindboggle extracts one fundus from each fold by finding the deepest vertices inside the fold, finding endpoints along the edge of the fold, connecting the former to the latter with tracks that run along deep and curved paths, and running a final filtration step. Just as anatomical labels segment folds into sulci, sulcus labels segment fold fundi into sulcal fundi.

##### Step 6: Segment cortical surface features with labels

- sulci from folds
- fundus per sulcus

Since folds are defined as deep, connected areas of a surface, and since folds may be connected to each other in ways that differ across brains, there usually does not exist a one-to-one mapping between folds of one brain and those of another. To address the correspondence problem, we need to find just those portions of the folds that correspond across brains. To accomplish this, Mindboggle segments folds into sulci, which do have a one-to-one correspondence across non-pathological brains (right side of Fig 7). Mindboggle defines a sulcus as a folded portion of cortex whose opposing banks are labeled with one or more sulcus label pairs in the DKT labeling protocol. Each label pair is unique to one sulcus and represents a boundary between two adjacent gyri, and each vertex has one gyrus label. The `extract_sulci` function assigns vertices in a fold to a sulcus in one of two cases. In the first case, if a vertex has a label that is in only one label pair in the fold, it is assigned that label pair’s sulcus if it can be connected through vertices with one of the pair’s labels to the boundary between the two labels. In the second case, the `segment_regions` function propagates labels from a label boundary to vertices whose labels are in multiple label pairs in the fold. Once sulci are defined, the `segment_by_region` function uses sulcus labels to segment fold fundi into sulcal fundi, which, like sulci, are features with one-to-one correspondence across non-pathological brains.

##### Step 7: Compute shape measures for each cortical surface label or sulcus

- surface area
- Laplace-Beltrami spectrum
- Zernike moments

In addition to shape measures computed for each vertex of a surface (Step 4), Mindboggle also computes shape measures that apply to collections of vertices such as gyri and sulci (Step 6): surface area (sum of surface areas across vertices), Laplace-Beltrami spectra, and Zernike moments.

Martin Reuter established important properties of the spectrum that relates to a shape’s intrinsic geometry with his “Shape-DNA” method [104–106]. This approach is specifically valuable for non-rigid shapes, such as anatomical structures: it is insensitive to local bending, as it quantifies only non-isometric deformation, e.g., stretching. The spectrum corresponds to the frequencies of the modes of the shape and its real-valued components, the eigenvalues, therefore describe different levels of detail (from more global low-frequency features to localized high-frequency details, Fig 9). The eigen-decomposition of the Laplace-Beltrami operator is computed via a finite element method (FEM). Mindboggle’s Python `fem_laplacian` function is based on Reuter’s Shape-DNA Matlab implementation, and their eigenvalues agree to the 16th decimal place, attributable to machine precision.

To calculate the distance between the descriptors of two shapes, Reuter describes several approaches, e.g., L^p^-norm, Hausdorff distance and weighted distances. One of the more prominent and simple distance measures is the Euclidean distance (L2 norm) of the first N smallest (non-zero) eigenvalues, where N is called the truncation parameter. To account for the linearly increasing magnitude of the eigenvalues (Weyl’s law), Reuter recommends to divide each value by its area and its index (done by default in Mindboggle). As an alternative, the Weighted Spectral Distance (WESD) [107] is included in Mindboggle (but not used by default). It computes the L^p^-norm of a weighted difference between the vectors of the N smallest Eigenvalues. This approach forms a pseudo-metric and also avoids domination of higher components on the final distance, making it insensitive to the truncation parameter N (with a decreasing influence as N gets larger). Additionally, the choice of p (for the L^p^-norm) influences how sensitive the metric is to finer as opposed to coarser differences in the shape; as p increases, WESD becomes less sensitive to differences at finer scales.

**Fig 9.**
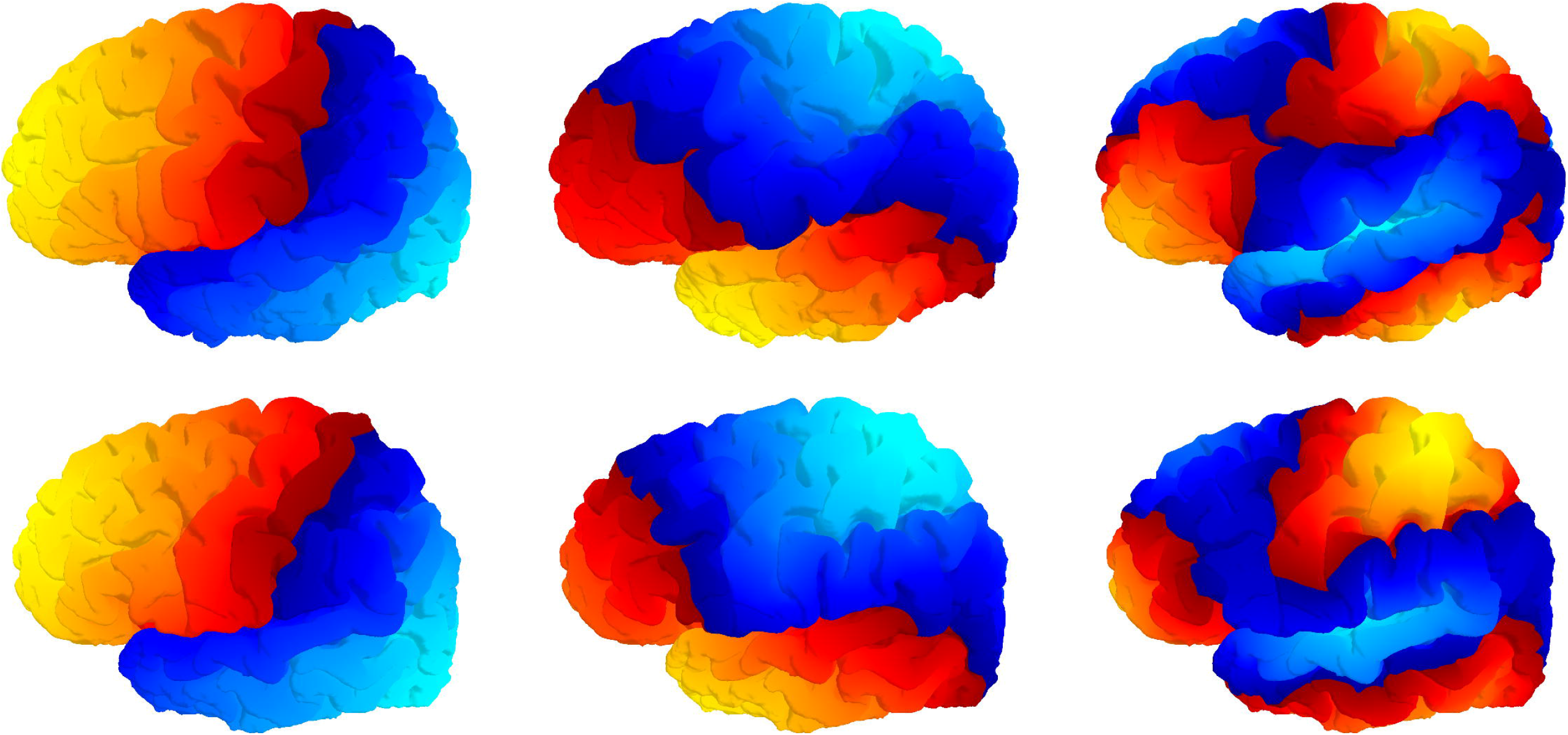
Laplace-Beltrami spectra. Mindboggle computes a Laplace-Beltrami spectrum for each feature (gyrus, sulcus, etc.), which relates to its intrinsic geometry, after Reuter et al.’s “Shape-DNA” method [104–106]. The components of the spectrum correspond roughly to the level of detail of the shape, from global to local, shown left to right for the 2nd, 3rd, and 9th spectral components for two different left brain hemispheres (top and bottom).

Moments can describe the shape of objects, images, or statistical distributions of points, and different types of moments confer different advantages [108]. Geometric moments of 3-D coordinates have been used to construct shape descriptors for human brain morphometry [109] because of desirable characteristics such as invariance to rotation, symmetry, and scale, and they can be computed for any topology. Zernike moments [110] have also been applied to human brain morphometry for classifying dementia patients [55] and confer several advantages over geometric moments. They form a set of orthogonal descriptors, where each descriptor contains independent information about the structure, allowing the original shape to be reconstructed from the moments. They have been extensively characterized for shape retrieval performance and are robust to noise. Zernike moments can also be calculated at different orders (levels of detail): low order moments represent low frequency information while high orders represent high frequency information. Mikhno et al. [55] implemented Pozo et al.’s [111] efficient 3-D implementation of Zernike moments in Matlab, and helped us test our Python implementation to ensure they give consistent results. The length of the descriptors exponentially increases with order, so order 20 yields 121 descriptors while order 35 yields 342, for example. Values are generally less than or equal to one, with values much greater than one indicating instability in the calculation, which could be due to the way the mesh is created or due to calculating at an order that is too high given the resolution or size of the object.

##### Step 8: Compute statistics for each shape measure in Step 4 for collections of vertices

- median
- median absolute deviation
- mean
- standard deviation
- skewness
- kurtosis
- lower quartile
- upper quartile

There can be thousands of vertices in a single feature such as a gyrus, sulcus, or fundus, so it makes sense to characterize a feature’s shape as either a distribution of per-vertex shape values (Step 4), or as a single shape value (Step 7). Mindboggle’s `stats_per_label` function generates tables containing both, with summary statistical measures representing the distributions of per-vertex shape values.

### 2.4. Mindboggle output

Example output data generated by Mindboggle is accessible at http://osf.io/8cf5z/. As with the input formats, volume files are in NIfTI format, surface meshes are in VTK format, and tables are comma-delimited. Each file contains integers that correspond to anatomical labels or features (0-24 for sulci). All output data are in the original subject’s space, except for additional surfaces and mean coordinates in MNI152 space [112]. The **Appendix** contains a directory tree with outputs from most of the optional arguments, and does not include interim results stored in a working directory or downloaded files in a cache directory.

## 3. Results

Mindboggle has been and continues to be subjected to a variety of evaluations (https://osf.io/x3up7/) and applied in a variety of contexts. In this section, we compare related shape measures (**3.1**), evaluate fundus extraction algorithms (**3.2**), and evaluate the consistency of shape measures between scans (**3.3**). We also demonstrate Mindboggle’s utility in measuring shape differences between left and right hemispheres (**3.4**), and in measuring brain shape variation (**3.5**).

### 3.1. Comparisons between brain shape measures

We compared shape measures with one another in a representative individual from the Mindboggle-101 data set (Fig 10) and for the entire data set (Fig 11, Fig 12, and Fig 13) to emphasize to the reader that shape measures are not independent of one another and that care must be taken when comparing differently defined shape measures or when using one as a proxy for another. Fig 10 plots over 130,000 vertices of one brain hemisphere, where the coordinates are two different shape measures assigned to each vertex: geodesic depth by travel depth (top) and mean curvature by travel depth (bottom). This figure demonstrates that curvature is positively correlated with depth and that geodesic depth produces higher shape values than travel depth, and may exaggerate depth, such as in the insula (also clearly evident in Fig 6 and Fig 11).

**Fig 10.**
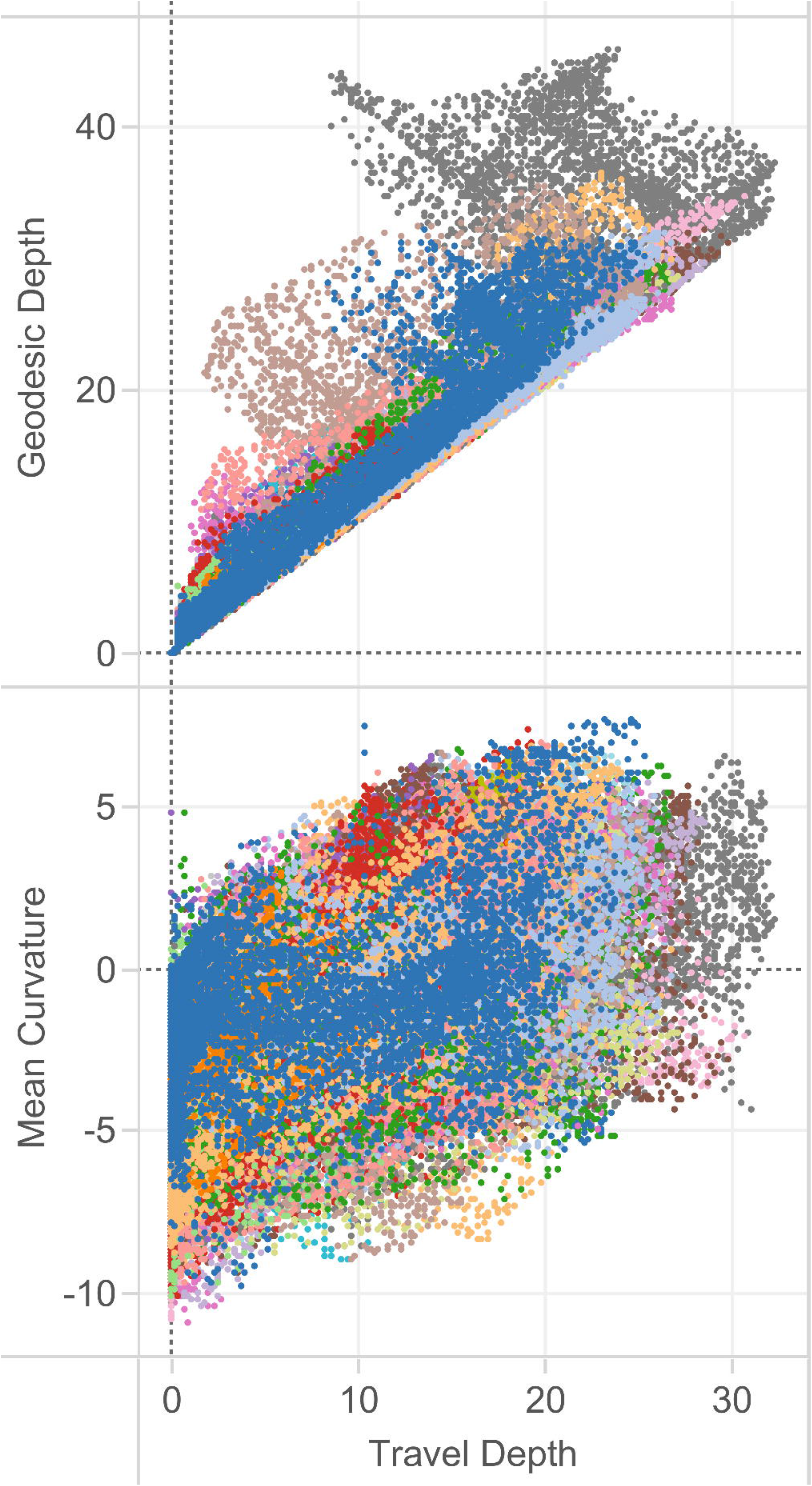
Relationships between brain shape measures. In these plots, we compare a pair of shape measures for each vertex of each right cortical region in a representative individual from the Mindboggle-101 brains, colored arbitrarily by region. Top: In this plot comparing two measures of depth, geodesic depth is higher than travel depth, and may exaggerate depth, such as in the insula (gray dots extending to the upper left). Bottom: In this plot of mean curvature by travel depth, we again see that the shape measures are not independent of one another. As one might expect, we see greater curvature at greater depth.

**Fig 11.**
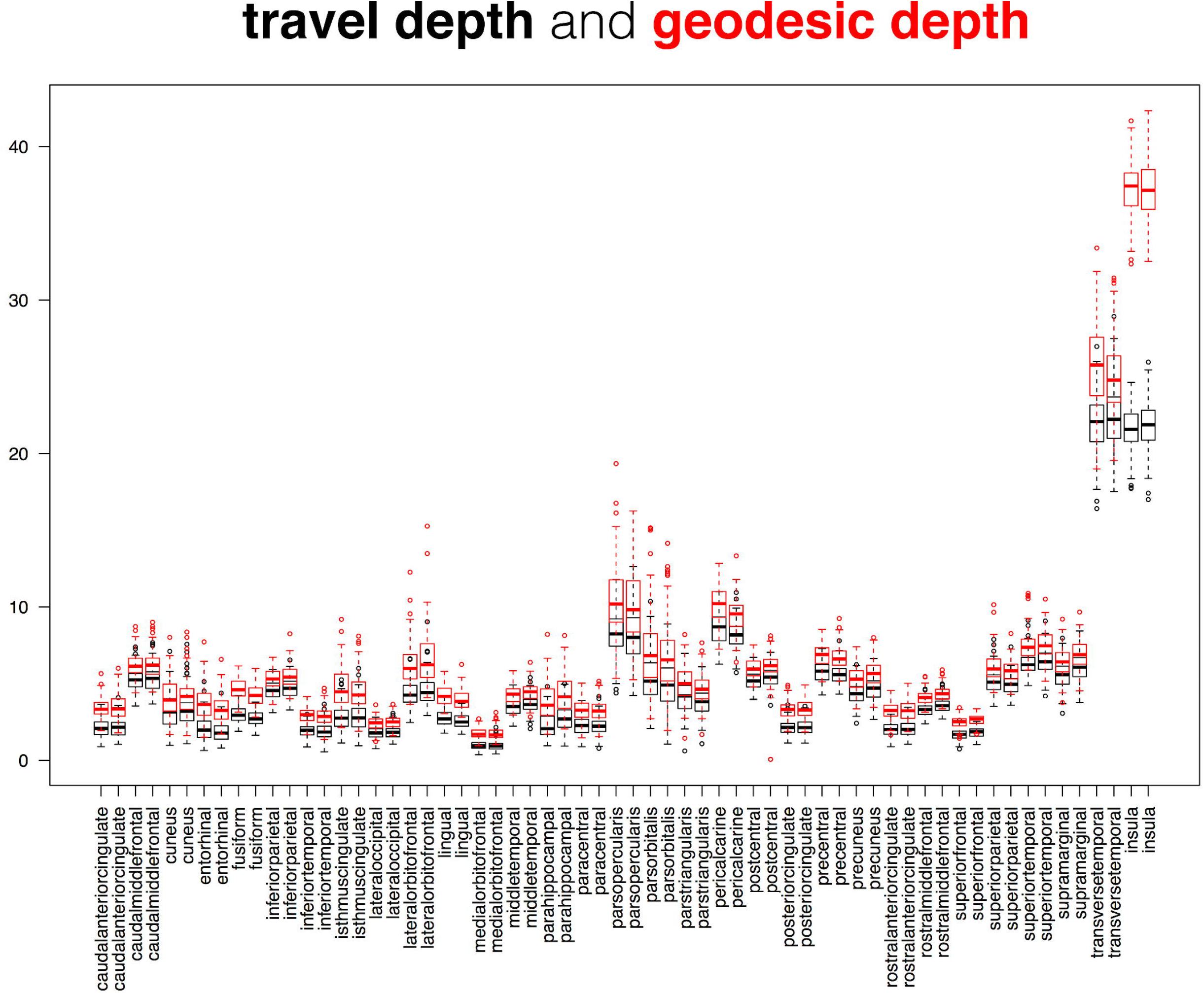
Comparison between cortical depth measures. This superposition of two box and whisker plots is a comparison between two measures of cortical surface depth applied to the 101 Mindboggle-101 brains: Mindboggle’s travel depth and geodesic depth. These surface measures are computed for every mesh vertex, so the plots were constructed from median depth values, with one value per labeled region. Geodesic depth deviates most from travel depth for the insular regions (far right).

**Fig 12.**
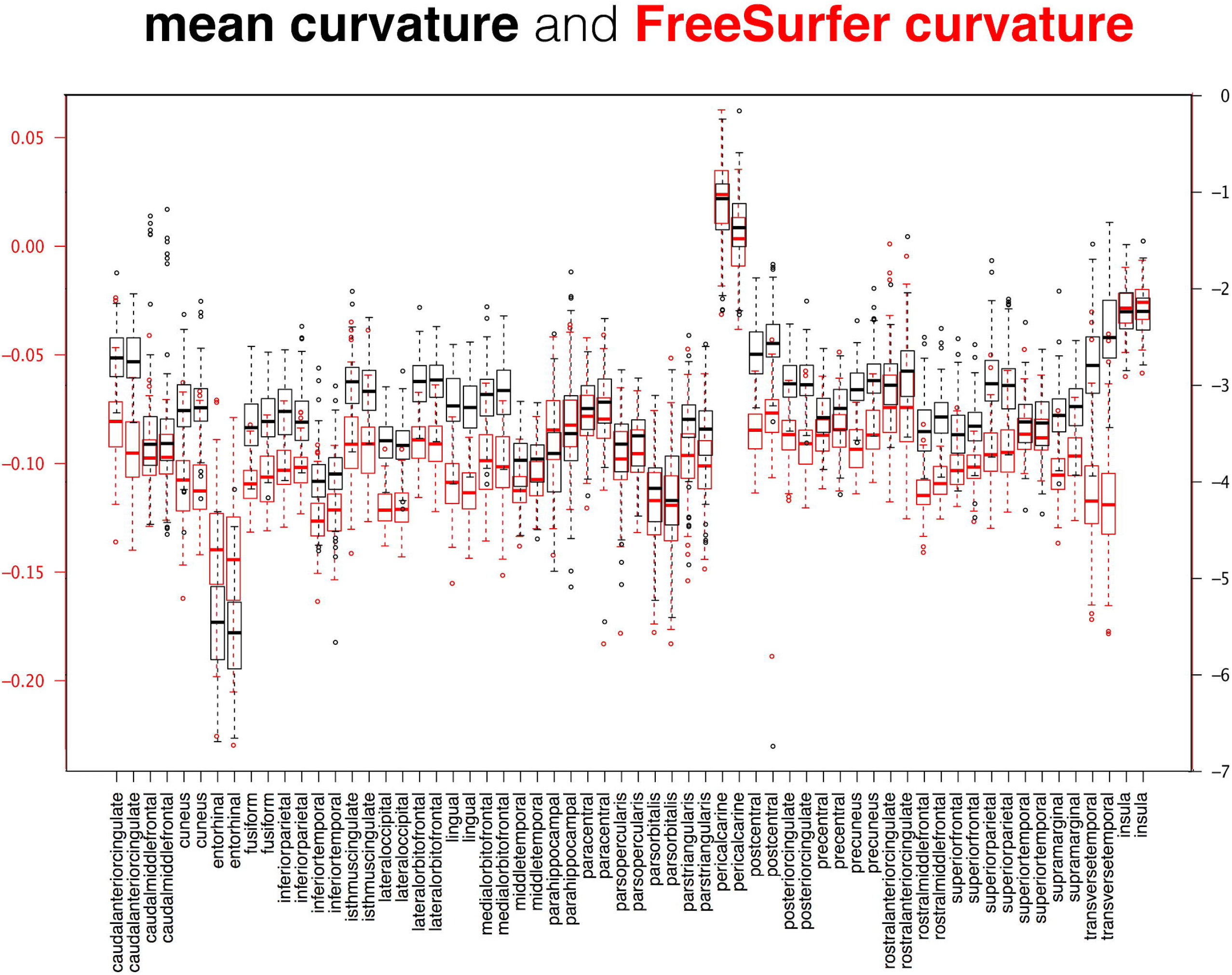
Comparison between cortical curvature measures. This superposition of two box and whisker plots is a comparison between two measures of cortical surface curvature applied to the 101 Mindboggle-101 brains: Mindboggle’s mean curvature and FreeSurfer’s curvature measure. These surface measures are computed for every mesh vertex, so the plots were constructed from median curvature values, with one value per labeled region. Deviations between the two curvature values are most evident for the entorhinal regions (fourth pair from the left).

**Fig 13.**
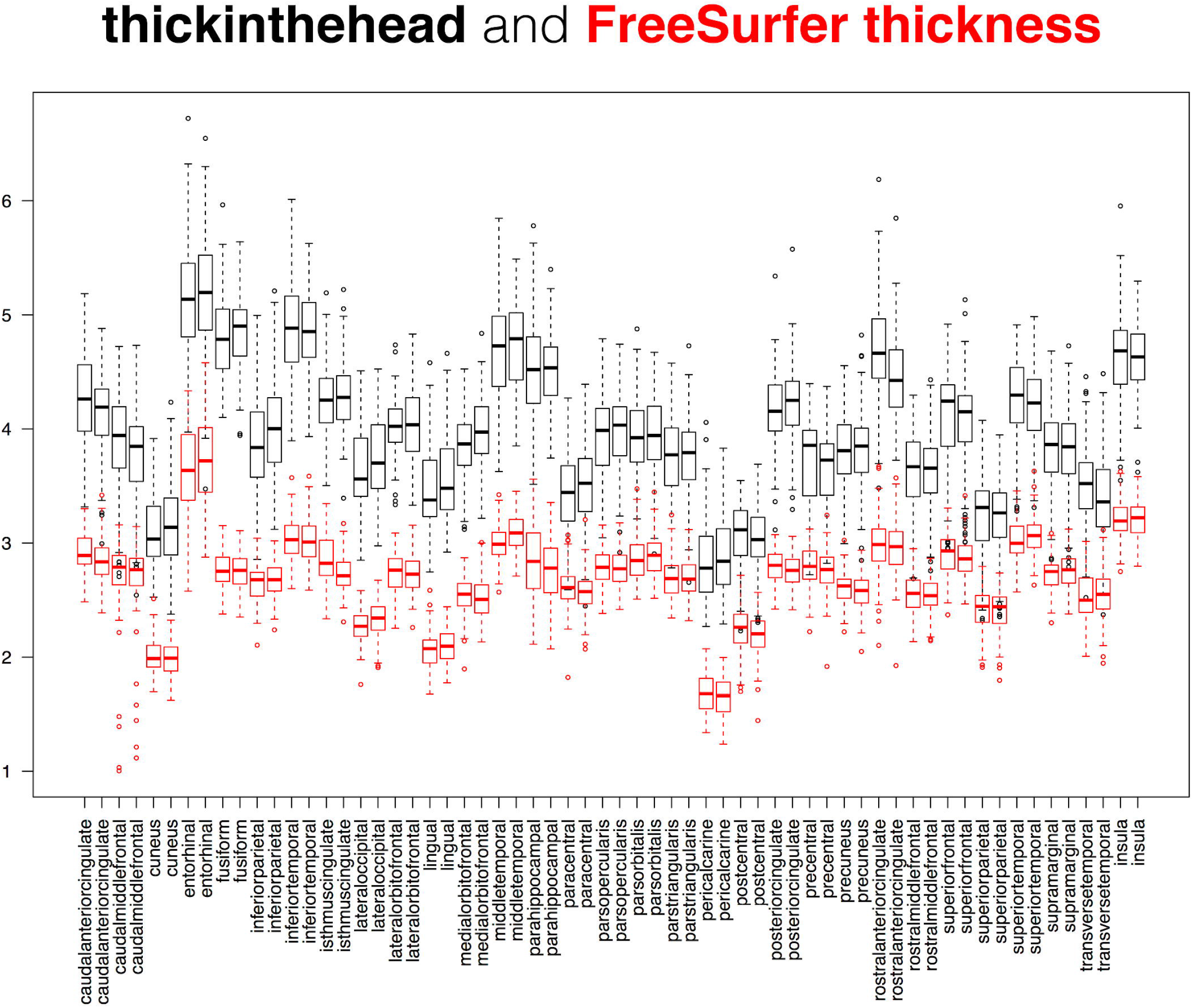
Comparison between cortical thickness measures. This superposition of two box and whisker plots is a comparison between two measures of cortical thickness applied to the 101 Mindboggle-101 brains: Mindboggle’s `thickinthehead` (black) and FreeSurfer’s thickness (red) measures. FreeSurfer’s thickness is defined per surface mesh vertex, so the red plot was constructed from median thickness values, with one value per labeled region.

While it may be useful to compare the distributions of two different shape measures for each region over a population (as in Fig 11, Fig 12, and Fig 13), we also computed the distance correlation between related shape measures for each cortical region in the Mindboggle-101 subjects (Table 1; https://osf.io/9cn7s/). To compare related (travel and geodesic depth, mean and FreeSurfer curvature) surface shape measures, we computed the distance correlation between each pair of shape measures across all of the vertices per region, and computed the average of the distance correlations per region across the 101 subjects. Distance correlation enabled a comparison of the pattern of values for a given region between two shape measures without regard for their absolute values. Mindboggle’s travel depth and geodesic depth measures were very highly correlated for 60 of the 62 regions, with distance correlations ranging from 0.91 to 1.00 (all but four greater than 0.95). The two outliers were the left and right insula (0.29 and 0.31), which corroborates our earlier assertion that geodesic depth can exaggerate depth values compared to travel depth in regions such as the insula. Mindboggle’s mean curvature and FreeSurfer’s curvature measures had distance correlations ranging from 0.73 (insula) to 0.91 (rostral middle frontal), with the top 10 values all for frontal and parietal regions. Since `thickinthehead` values are computed per region, not per vertex, to compare `thickinthehead` with median FreeSurfer thickness values, we constructed a pair of vectors for each region with 101 values, each value corresponding to the shape measure for that region in a subject, and computed the distance correlation between the two vectors. The highest distance correlations (0.8 to 0.7) were obtained by frontal and parietal regions, and the lowest correlations (0.3 to 0.2) by precuneus, parahippocampal, fusiform, and cingulate regions.

**Table 1.** Distance correlations between related shape measures. To compare pairs of related (travel and geodesic depth, mean and FreeSurfer curvature) surface shape measures, we computed the distance correlation between vectors of shape values for all vertices in each cortical region, and averaged the distance correlations across the 101 Mindboggle-101 subjects. For `thickinthehead` and FreeSurfer thickness measures, we computed the distance correlation between vectors of median shape values for all 101 Mindboggle-101 subjects for each cortical region.

| Cortical region | travel depth vs.<br>geodesic depth |  | mean curvature vs.<br>FreeSurfer curvature |  | thickinthehead vs.<br>FreeSurfer thickness |  |
| --- | --- | --- | --- | --- | --- | --- |
|  | left | right | left | right | left | right |
| caudal anterior cingulate | 0.97 | 0.96 | 0.83 | 0.83 | 0.39 | 0.38 |
| caudal middle frontal | 0.99 | 0.99 | 0.88 | 0.88 | 0.72 | 0.70 |
| cuneus | 0.99 | 0.99 | 0.84 | 0.83 | 0.52 | 0.51 |
| entorhinal | 0.96 | 0.96 | 0.77 | 0.75 | 0.38 | 0.33 |
| fusiform | 0.97 | 0.97 | 0.83 | 0.83 | 0.22 | 0.20 |
| inferior parietal | 0.99 | 0.99 | 0.88 | 0.88 | 0.56 | 0.50 |
| inferior temporal | 0.98 | 0.98 | 0.84 | 0.84 | 0.31 | 0.39 |
| isthmus cingulate | 0.91 | 0.93 | 0.78 | 0.79 | 0.19 | 0.30 |
| lateral occipital | 0.99 | 0.99 | 0.86 | 0.86 | 0.54 | 0.57 |
| lateral orbitofrontal | 0.92 | 0.92 | 0.80 | 0.81 | 0.54 | 0.54 |
| lingual | 0.97 | 0.98 | 0.83 | 0.82 | 0.45 | 0.62 |
| medial orbitofrontal | 0.97 | 0.97 | 0.82 | 0.82 | 0.42 | 0.57 |
| middle temporal | 0.99 | 1.00 | 0.88 | 0.88 | 0.49 | 0.40 |
| parahippocampal | 0.96 | 0.97 | 0.81 | 0.84 | 0.44 | 0.26 |
| paracentral | 0.99 | 0.99 | 0.87 | 0.87 | 0.64 | 0.59 |
| pars opercularis | 0.98 | 0.98 | 0.89 | 0.89 | 0.65 | 0.47 |
| pars orbitalis | 0.98 | 0.98 | 0.90 | 0.90 | 0.43 | 0.50 |
| pars triangularis | 1.00 | 1.00 | 0.90 | 0.90 | 0.63 | 0.47 |
| pericalcarine | 0.96 | 0.97 | 0.76 | 0.78 | 0.34 | 0.37 |
| postcentral | 1.00 | 1.00 | 0.87 | 0.87 | 0.71 | 0.63 |
| posterior cingulate | 0.99 | 0.99 | 0.84 | 0.83 | 0.29 | 0.38 |
| precentral | 0.99 | 0.99 | 0.88 | 0.88 | 0.70 | 0.54 |
| precuneus | 0.98 | 0.98 | 0.86 | 0.86 | 0.29 | 0.48 |
| rostral anterior cingulate | 0.98 | 0.97 | 0.80 | 0.79 | 0.26 | 0.35 |
| rostral middle frontal | 0.99 | 0.99 | 0.91 | 0.91 | 0.75 | 0.61 |
| superior frontal | 0.99 | 0.99 | 0.89 | 0.89 | 0.80 | 0.71 |
| superior parietal | 0.99 | 0.99 | 0.89 | 0.89 | 0.69 | 0.76 |
| superior temporal | 0.99 | 0.99 | 0.86 | 0.85 | 0.59 | 0.52 |
| supramarginal | 1.00 | 1.00 | 0.88 | 0.87 | 0.65 | 0.60 |
| transverse temporal | 0.97 | 0.98 | 0.76 | 0.74 | 0.67 | 0.69 |
| insula | 0.29 | 0.31 | 0.73 | 0.73 | 0.46 | 0.38 |

#### 3.1.1. Comparison between travel depth and FreeSurfer’s convexity measure

As described above, travel depth uses a reference wrapper surface that lies closer to the cortical surface than a convex hull would. In particular, the wrapper lies closer to the medial temporal lobe, so the gyri in this area have depth values equal to zero as one would want. FreeSurfer’s convexity measure [80], often used to indicate relative depth, leads to non-zero and even negative values for vertices on these gyri (Fig 4 in **Supplement 1**). We computed the mean and standard deviation of four statistical measures of travel depth and FreeSurfer’s convexity values for over 130,000 vertices in a representative cortical surface. For this comparison, we consider a point to be close to the wrapper surface if the distance between the two is smaller than 0.1 mm, a depth value is considered small if it is less than 0.1 mm, and a convexity value is considered small if it is less than the smallest convexity value for all the vertices in the mesh. For travel depth, by definition all vertices (and only those vertices) that are close to the wrapper surface have a small depth. For convexity, almost all vertices (97.71%) that have a small convexity value are close to the wrapper surface, but they represent only 6.89% of the vertices close to the wrapper surface (Table 1 in **Supplement 1**). One conclusion we drew from this comparison is that while both travel depth and FreeSurfer’s convexity measures represent depth well for deep portions of a surface, travel depth provides a more faithful representation for shallow portions.

#### 3.1.2. Comparison between cortical thickness measures

We are aware of only one study directly comparing FreeSurfer with manual cortical thickness measures, where the manual estimates were made in nine gyral crowns of a post-mortem brain, selected for their low curvature and high probability of having been sampled perpendicular to the plane of section [113]. We compared `thickinthehead`, FreeSurfer, and ANTs cortical thickness estimates in different populations, including the Mindboggle-101 subjects (Fig 13) and in the 40 EMBARC control subjects (https://osf.io/jwhea/). For 16 cortical regions in the 40 subjects, we measured scan-rescan reliability of cortical thickness measures, and we compared thickness measures with published estimates based on manual delineations of MR images of living brains [114]. Forty percent of FreeSurfer estimates for the labels were in the published ranges of values, whereas almost ninety percent of `thickinthehead`’s estimates were within these ranges (as mentioned above, Klauschen observed that FreeSurfer underestimates gray matter and overestimates white matter [77]). ANTs values deviated further from the published estimates and were less reliable (greater inter-scan and inter-subject ranges) than the FreeSurfer or `thickinthehead` values.

### 3.2. Evaluation of fundus extraction algorithms

This section presents the first quantitative comparison of fundus extraction software algorithms. Since there exists no ground truth for fundus curves, we must resort to other means of evaluation. We leave it to future work to determine their utility for practical applications such as diagnosis and prediction of disorders. Since the DKT labeling protocol defines many of its anatomical label boundaries along approximations of fundus curves, we used the manually edited anatomical label boundaries in the Mindboggle-101 dataset as gold standard data to evaluate the positions of fundi extracted by four different algorithms in 2013. Specifically, for each of the 48 fundi/sulci defined by the DKT protocol, we computed the mean of the minimum Euclidean distances from the label boundary vertices in the sulcus to the fundus vertices in the sulcus, as well as from the fundus vertices in the sulcus to the label boundary vertices in the sulcus. The algorithms included Mindboggle’s default `connect_points_erosion` function described above, Forrest Bao’s pruned minimum spanning tree algorithm [102], Gang Li’s algorithm [115], and an algorithm in the BrainVisa software [95]. The final algorithm was omitted from the results because too few fundi were extracted to make an adequate comparison (BrainVisa extracts 65 sulci per hemisphere, and it is possible that the program did not define some folds as sulci that contain fundi according to the DKT labeling protocol).

All of the fundi, summary statistics, and results are available online (https://osf.io/r95wb/). While there was no clear winner, we can summarize our comparison by computing the mean distance between fundi and label boundaries across all sulci for the three methods and by tallying how many sulci had the smallest mean distance among the methods. When measured from label boundaries to fundi, Gang Li’s and Mindboggle’s fundi were closer than were Forrest Bao’s (mean distances of 2.09mm and 2.38mm vs. 3.65mm, respectively; 25 and 21 vs. 2 closest sulci), whereas when measured from fundi to label boundaries, Forrest Bao’s fundi were closer than were Mindboggle’s or Gang Li’s (mean distances of 3.33mm vs. 4.06mm and 4.65mm, respectively; 41 vs. 5 and 2 closest sulci). When measuring from either direction, the maximum distances averaged across all sulci were higher for Forrest Bao’s fundi (11.65mm and 11.61mm) than for Mindboggle’s (10.84mm and 9.75mm) or Gang Li’s (11.12mm and 6.87mm).

### 3.3. Consistency of shape measures between MRI scans of the same person

For a shape measure to be useful in comparative morphometry, it should be more sensitive to differences in anatomy than to differences in MRI scanning setup or artifacts. To get a sense of the degree of scan/rescan consistency of our shape measures, we ran Mindboggle on 41 Mindboggle-101 subjects with a second MRI scan (OASIS-TRT-20 and MMRR-21 cohorts). We computed the fractional shape difference per cortical region as the absolute value of the difference between the region’s shape values for the two scans divided by the first scan’s shape value. For the volumetric shape measures (volume and `thickinthehead` cortical thickness), shape value is computed by region; for the surface-based shape measures (area, travel and geodesic depth, mean and FreeSurfer curvature, and FreeSurfer thickness), shape value is assigned the median value across all vertices within a region. All shape tables, statistical summary tables, and accompanying plots are available online (https://osf.io/mhc37/).

Table 2a gives the average across the 41 subjects of the fractional shape differences between MRI scans for each of the 31 left cortical regions, and for each shape measure, and Table 2b gives a statistical summary of the differences. In general, the values are low enough to suggest high inter-scan shape consistency, but we will point out values greater than or equal to 0.10. Of the volumetric shape measures (volume and `thickinthehead`), only one value exceeded or equaled 0.10: entorhinal volume (0.21). Entorhinal cortex had the second smallest volume of manually labeled MRI cortical regions in 101 healthy human brains [22] (after transverse temporal cortex; see https://osf.io/st7nk/), and low scan/rescan consistency for small brain structures corroborates Jovicich’s observation in 2013 [116]: “We found that the smaller structures (pallidum and amygdala) yielded the highest absolute volume reproducibility errors, approximately 3.8% (average across sites), whereas all other structures had errors in the range 1.8-2.2% (average across sites), with the longitudinal segmentation analysis. Our absolute % errors in test-retest volumetric estimates are comparable to those reported by previous studies (Kruggel et al., 2010; Morey et al., 2010; Reuter et al., 2012).” Regarding cortical thickness measures, Jovicich observed: “The thickness reproducibility results of the various structures were largely consistent across sites and vendors, with errors in the range 0.8 – 5.0% for the longitudinal analysis (Table 7).” Of the surface shape measures, the following exceeded or equaled 0.10 for three measures (travel depth, geodesic depth, and FreeSurfer curvature): entorhinal, medial orbitofrontal, and (caudal anterior, rostral anterior, and isthmus) cingulate regions; and for at least one of the measures: lateral orbitofrontal, parahippocampal, pericalcarine, and insular regions. The greatest differences were for FreeSurfer curvature in the pericalcarine (0.34), insula (0.28), and rostral anterior cingulate (0.23), followed by entorhinal volume (0.21) and travel depth (0.20). FreeSurfer curvature had the greatest number of outliers (Table 2b) and was the only shape measure that spanned negative to positive values, so regions with very small median curvature values could have inflated these fractions. Future evaluations will assess the impact that differences in scans have on morphometry-based clinical research.

**Table 2a.** Shape differences between MRI scans. This table lists shape differences between two scans of the same brain averaged across 41 brains. The shape differences are computed for each of the 31 left cortical regions as the absolute value of the difference between the region’s shape values between the two scans divided by the first scan’s shape value. For the surface-based shape values, we used the median value for all vertices within each region. [thick = `thickinthehead` cortical thickness; travel = travel depth; geodesic = geodesic depth; curv = mean curvature; FScurv = FreeSurfer’s curvature; FSthick = FreeSurfer’s thickness]

| **Cortical region** | **volume** | **thick** | **area** | **travel** | **geodesic** | **curv** | **FScurv** | **FSthick** |
| --- | --- | --- | --- | --- | --- | --- | --- | --- |
| caudal anterior cingulate | 0.04 | 0.05 | 0.05 | 0.10 | 0.10 | 0.06 | 0.18 | 0.04 |
| caudal middle frontal | 0.04 | 0.05 | 0.05 | 0.03 | 0.03 | 0.05 | 0.07 | 0.04 |
| cuneus | 0.04 | 0.06 | 0.04 | 0.06 | 0.05 | 0.03 | 0.05 | 0.05 |
| entorhinal | 0.21 | 0.06 | 0.09 | 0.20 | 0.16 | 0.06 | 0.14 | 0.05 |
| fusiform | 0.03 | 0.04 | 0.03 | 0.04 | 0.05 | 0.03 | 0.05 | 0.03 |
| inferior parietal | 0.03 | 0.04 | 0.02 | 0.03 | 0.02 | 0.04 | 0.05 | 0.04 |
| inferior temporal | 0.05 | 0.04 | 0.03 | 0.07 | 0.06 | 0.03 | 0.06 | 0.02 |
| isthmus cingulate | 0.04 | 0.08 | 0.07 | 0.11 | 0.13 | 0.04 | 0.11 | 0.04 |
| lateral occipital | 0.03 | 0.02 | 0.03 | 0.08 | 0.07 | 0.03 | 0.04 | 0.04 |
| lateral orbitofrontal | 0.09 | 0.05 | 0.04 | 0.08 | 0.10 | 0.05 | 0.09 | 0.04 |
| lingual | 0.03 | 0.04 | 0.04 | 0.05 | 0.06 | 0.03 | 0.04 | 0.04 |
| medial orbitofrontal | 0.07 | 0.06 | 0.06 | 0.14 | 0.10 | 0.06 | 0.11 | 0.05 |
| middle temporal | 0.03 | 0.05 | 0.03 | 0.04 | 0.03 | 0.03 | 0.05 | 0.03 |
| parahippocampal | 0.04 | 0.04 | 0.04 | 0.11 | 0.12 | 0.06 | 0.09 | 0.03 |
| paracentral | 0.07 | 0.04 | 0.05 | 0.08 | 0.08 | 0.04 | 0.07 | 0.06 |
| pars opercularis | 0.04 | 0.06 | 0.04 | 0.03 | 0.03 | 0.03 | 0.06 | 0.03 |
| pars orbitalis | 0.06 | 0.05 | 0.04 | 0.06 | 0.07 | 0.05 | 0.09 | 0.04 |
| pars triangularis | 0.03 | 0.06 | 0.03 | 0.06 | 0.05 | 0.04 | 0.09 | 0.04 |
| pericalcarine | 0.06 | 0.06 | 0.04 | 0.03 | 0.03 | 0.10 | 0.34 | 0.07 |
| postcentral | 0.04 | 0.02 | 0.03 | 0.03 | 0.03 | 0.03 | 0.05 | 0.06 |
| posterior cingulate | 0.04 | 0.08 | 0.05 | 0.09 | 0.09 | 0.05 | 0.09 | 0.04 |
| precentral | 0.04 | 0.03 | 0.02 | 0.02 | 0.02 | 0.03 | 0.04 | 0.04 |
| precuneus | 0.03 | 0.05 | 0.03 | 0.03 | 0.04 | 0.03 | 0.07 | 0.04 |
| rostral anterior cingulate | 0.04 | 0.09 | 0.06 | 0.10 | 0.10 | 0.08 | 0.23 | 0.05 |
| rostral middle frontal | 0.04 | 0.07 | 0.03 | 0.04 | 0.03 | 0.05 | 0.06 | 0.04 |
| superior frontal | 0.02 | 0.04 | 0.02 | 0.06 | 0.04 | 0.03 | 0.04 | 0.03 |
| superior parietal | 0.04 | 0.03 | 0.02 | 0.03 | 0.03 | 0.04 | 0.05 | 0.05 |
| superior temporal | 0.04 | 0.05 | 0.03 | 0.02 | 0.03 | 0.02 | 0.04 | 0.02 |
| supramarginal | 0.04 | 0.04 | 0.03 | 0.04 | 0.04 | 0.04 | 0.07 | 0.04 |
| transverse temporal | 0.06 | 0.07 | 0.04 | 0.02 | 0.03 | 0.04 | 0.07 | 0.06 |
| insula | 0.03 | 0.09 | 0.03 | 0.01 | 0.01 | 0.05 | 0.28 | 0.03 |

**Table 2b.** Summary statistics of shape differences between MRI scans. This table gives a statistical summary of the shape differences between two scans of the same brain for 41 brains. The “mean” column is the average of the mean values in Table 1a, while the other columns contain averages of their respective values over the 31 regions; for example, the “std” column contains the average of the std values computed for each of the 31 regions. [>0.50 and >0.25 give the number of regions (out of 1,271 = 31 regions times 41 subjects) where the fractional absolute difference was above 0.50 and 0.25, respectively.]

|  | mean | std | min | 25% | 50% | 75% | max | >0.50 | >0.25 |
| --- | --- | --- | --- | --- | --- | --- | --- | --- | --- |
| volume | 0.047 | 0.044 | 0.002 | 0.019 | 0.035 | 0.063 | 0.213 | 5 | 21 |
| thickinthehead | 0.052 | 0.046 | 0.001 | 0.017 | 0.039 | 0.078 | 0.183 | 0 | 16 |
| area | 0.039 | 0.038 | 0.001 | 0.014 | 0.030 | 0.054 | 0.182 | 2 | 6 |
| travel depth | 0.061 | 0.050 | 0.002 | 0.024 | 0.050 | 0.085 | 0.229 | 1 | 36 |
| geodesic depth | 0.059 | 0.049 | 0.002 | 0.022 | 0.048 | 0.082 | 0.222 | 2 | 35 |
| mean curvatures | 0.044 | 0.038 | 0.001 | 0.016 | 0.033 | 0.059 | 0.170 | 0 | 7 |
| FreeSurfer curvature | 0.094 | 0.091 | 0.004 | 0.033 | 0.070 | 0.127 | 0.433 | 17 | 78 |
| FreeSurfer thickness | 0.041 | 0.036 | 0.001 | 0.014 | 0.032 | 0.060 | 0.147 | 0 | 0 |

### 3.4. Measuring shape differences between left and right hemispheres

To measure interhemispheric shapes differences, we computed the fractional shape difference per cortical region as in the preceding section, replacing inter-scan differences with interhemispheric differences (https://osf.io/dp4zy/), and using all 101 Mindboggle-101 brains. Table 3a gives the average across the 101 subjects of the fractional shape differences between hemispheres for each of the 31 cortical regions, and for each shape measure, and Table 3b gives a statistical summary of the differences. The values are much higher than the corresponding inter-scan differences in the previous section, suggesting that shape differences between hemispheres are greater than shape differences between MRI scans of the same hemisphere.

**Table 3a.** Shape differences between left and right hemispheres. Shape differences between hemispheres are computed for each of the 31 cortical regions in all 101 of the Mindboggle-101 subjects as the absolute value of the difference between the region’s left and right shape values divided by the left shape value. For the surface-based shape values, we used the median value for all vertices within each region. (Refer to Table 1a caption for abbreviations.)

| Cortical regions | volume | thick | area | travel | geodesic | curv | FScurv | FSthick |
| --- | --- | --- | --- | --- | --- | --- | --- | --- |
| caudal anterior cingulate | 0.26 | 0.06 | 0.32 | 0.27 | 0.21 | 0.11 | 0.32 | 0.06 |
| caudal middle frontal | 0.14 | 0.04 | 0.20 | 0.14 | 0.13 | 0.10 | 0.16 | inf |
| cuneus | 0.12 | 0.06 | 0.17 | 0.32 | 0.24 | 0.10 | 0.15 | 0.04 |
| entorhinal | 0.17 | 0.08 | 0.23 | 0.41 | 0.30 | 0.10 | 0.21 | 0.09 |
| fusiform | 0.08 | 0.05 | 0.15 | 0.14 | 0.13 | 0.07 | 0.10 | 0.04 |
| inferior parietal | 0.23 | 0.05 | 0.25 | 0.11 | 0.11 | 0.06 | 0.09 | 0.03 |
| inferior temporal | 0.09 | 0.04 | 0.14 | 0.21 | 0.16 | 0.07 | 0.08 | 0.05 |
| isthmus cingulate | 0.14 | 0.04 | 0.17 | 0.21 | 0.20 | 0.10 | 0.18 | 0.07 |
| lateral occipital | 0.09 | 0.04 | 0.14 | 0.22 | 0.18 | 0.05 | 0.07 | 0.04 |
| lateral orbitofrontal | 0.06 | 0.05 | 0.11 | 0.23 | 0.26 | 0.09 | 0.12 | 0.05 |
| lingual | 0.11 | 0.04 | 0.12 | 0.15 | 0.13 | 0.08 | 0.11 | 0.05 |
| medial orbitofrontal | 0.08 | 0.05 | 0.18 | 0.27 | 0.19 | 0.10 | 0.17 | 0.06 |
| middle temporal | 0.11 | 0.04 | 0.12 | 0.17 | 0.15 | 0.07 | 0.10 | 0.05 |
| parahippocampal | 0.11 | 0.05 | 0.13 | 0.36 | 0.29 | 0.15 | 0.19 | 0.08 |
| paracentral | 0.16 | 0.04 | 0.18 | 0.25 | 0.20 | 0.08 | 0.16 | 0.04 |
| pars opercularis | 0.18 | 0.04 | 0.33 | 0.20 | 0.23 | 0.10 | 0.17 | 0.05 |
| pars orbitalis | 0.22 | 0.05 | 0.32 | 0.37 | 0.40 | 0.13 | 0.20 | 0.06 |
| pars triangularis | 0.20 | 0.05 | 0.32 | 0.30 | 0.24 | 0.12 | 0.21 | 0.06 |
| pericalcarine | 0.18 | 0.06 | 0.21 | 0.10 | 0.11 | 0.45 | inf | 0.06 |
| postcentral | 0.09 | 0.03 | 0.12 | 0.13 | 0.13 | 0.10 | 0.14 | 0.05 |
| posterior cingulate | 0.13 | 0.04 | 0.13 | 0.21 | 0.18 | 0.09 | 0.15 | 0.05 |
| precentral | 0.07 | 0.03 | 0.09 | 0.10 | 0.10 | 0.07 | 0.11 | 0.04 |
| precuneus | 0.06 | 0.03 | 0.11 | 0.17 | 0.14 | 0.07 | 0.13 | 0.03 |
| rostral anterior cingulate | 0.22 | 0.07 | 0.36 | 0.21 | 0.18 | 0.10 | 1.23 | 0.06 |
| rostral middle frontal | 0.08 | 0.03 | 0.17 | 0.14 | 0.12 | 0.08 | 0.09 | 0.04 |
| superior frontal | 0.06 | 0.03 | 0.14 | 0.19 | 0.14 | 0.06 | 0.07 | 0.04 |
| superior parietal | 0.07 | 0.02 | 0.15 | 0.12 | 0.12 | 0.07 | 0.12 | 0.03 |
| superior temporal | 0.07 | 0.03 | 0.12 | 0.10 | 0.10 | 0.09 | 0.11 | 0.04 |
| supramarginal | 0.12 | 0.03 | 0.19 | 0.17 | 0.16 | 0.08 | 0.13 | 0.04 |
| transverse temporal | 0.24 | 0.06 | 0.24 | 0.10 | 0.10 | 0.18 | 0.27 | 0.06 |
| insula | 0.06 | 0.04 | 0.07 | 0.05 | 0.03 | 0.10 | 0.32 | 0.05 |

**Table 3b.** Summary statistics of shape differences between left and right hemispheres. This table gives a statistical summary of the interhemispheric shape differences for the 101 Mindboggle-101 brains. The “mean” column is the average of the mean values in Table 3a, while the other columns contain averages of their respective values over the 31 regions; for example, the “std” column contains the average of the std values computed for each of the 31 regions. [>0.50 and >0.25 give the number of regions (out of 3,131 = 31 regions times 101 subjects) where the fractional absolute difference was above 0.50 and 0.25, respectively.]

|  | mean | std | min | 25% | 50% | 75% | max | >0.50 | >0.25 |
| --- | --- | --- | --- | --- | --- | --- | --- | --- | --- |
| volume | 0.129 | 0.091 | 0.002 | 0.058 | 0.117 | 0.180 | 0.448 | 38 | 443 |
| thickinthehead | 0.044 | 0.036 | 0.001 | 0.017 | 0.037 | 0.064 | 0.169 | 0 | 4 |
| area | 0.183 | 0.163 | 0.002 | 0.074 | 0.148 | 0.248 | 1.025 | 165 | 744 |
| travel depth | 0.198 | 0.199 | 0.003 | 0.074 | 0.150 | 0.257 | 1.251 | 211 | 764 |
| geodesic depth | 0.173 | 0.163 | 0.002 | 0.067 | 0.133 | 0.229 | 1.009 | 148 | 658 |
| mean curvatures | 0.104 | 0.113 | 0.002 | 0.038 | 0.079 | 0.141 | 0.872 | 59 | 192 |
| FreeSurfer curvature | 0.190 | 0.435 | 0.003 | 0.074 | 0.150 | 0.250 | 3.890 | 205 | 626 |
| FreeSurfer thickness | 0.050 | 0.077 | 0.001 | 0.018 | 0.036 | 0.062 | 0.691 | 23 | 28 |

### 3.7. Measuring human brain shape variation

To estimate the normal range of variation in the shapes of healthy adult human brains, we applied Mindboggle software in 2015 to compute shape measures for our Mindboggle-101 dataset (see **2.1**: 2010 above). The result is the largest set of shape measures computed on healthy human brain data (See **Supplement 2** and https://osf.io/gzshf/ for detailed results) [117, 118]. We are treating these as normative data against which anyone can compare similarly processed images of different healthy populations as well as patient populations.

The data we analysed consist of repeated measurements on five distinct real-valued shape measures (mean curvature, geodesic depth, travel depth, FreeSurfer convexity, and FreeSurfer thickness) for each of 31 distinct regions per brain hemisphere in each of the 101 subjects. Each subject was scanned at one of five different laboratories. At the bottom of Fig 14 is one example of the many heatmap tables we have generated from these data (all results are accessible at https://osf.io/d7hx8/). Each table presents one value for each labeled region or sulcus for each of the 101 subjects. The value is either volume or `thickinthehead` cortical thickness for volumetric images, or for one of the five surface shape measures above, one of eight summary statistical measures (mean, median, median absolute deviation, standard deviation, lower and upper quartiles, skewness, and kurtosis) computed across all vertices in the surface mesh of the labeled region or sulcus.

We organized the data in a nested fashion: brain hemisphere is nested within subject, and subject is nested within laboratory. In addition to the five shape measurements and the three nested classification factors, the data also include three covariates: sex (male, female), age (integer variable), and handedness (left, right; we relabeled two ambidextrous subjects as left-handed). Given the grouped nature of the data, we used linear mixed models for the statistical modeling of the data. To assess the importance of each of the covariates and nested classification factors, we fitted 24 distinct linear mixed models for each shape measure and brain region combination to assess the importance of each of the covariates (sex, handedness, and age as fixed effects) and nested classification factors (laboratory, subject, and brain hemisphere as random effects). For each shape measure, we decomposed the total variance into the variance between laboratories, between subjects within a laboratory, between brain hemispheres within a subject, and within brain hemispheres.

For each shape measure and brain region combination, we used the Bayesian Information Criterion (BIC) score to select the best model among the 24 competing models. A BIC score is a goodness of fit measure used to perform model selection among models with different dimensions (number of parameters), and is proportional to the negative log likelihood of the model penalized by the number of parameters in the model. It strikes a balance between model fit (measured by the log-likelihood score) and model complexity (measured by the number of parameters in the model). In the context of linear models, an over-parameterized model will always have a larger log-likelihood score than a more parsimonious model, but it will also likely overfit the data. Nonetheless, by including a penalty proportional to the number of parameters in the model, the BIC score can be used to compare models with different dimensions since over-parameterized models are penalized to a greater extent. The smaller the BIC score, the better the model fits the data.

Two models stood out as the best models for the mean curvature, travel depth, FreeSurfer convexity, and FreeSurfer thickness shape measures across the 31 brain regions (**Supplement 2**). Both models include handedness and age as fixed effects. They only differ by the inclusion of the extra “subject within lab” nesting level. For all shape measures and brain regions, the bulk of the variability was concentrated in the residual, not in the hemisphere (“side”), subject, or laboratory (top of Fig 14).

**Fig 14.**
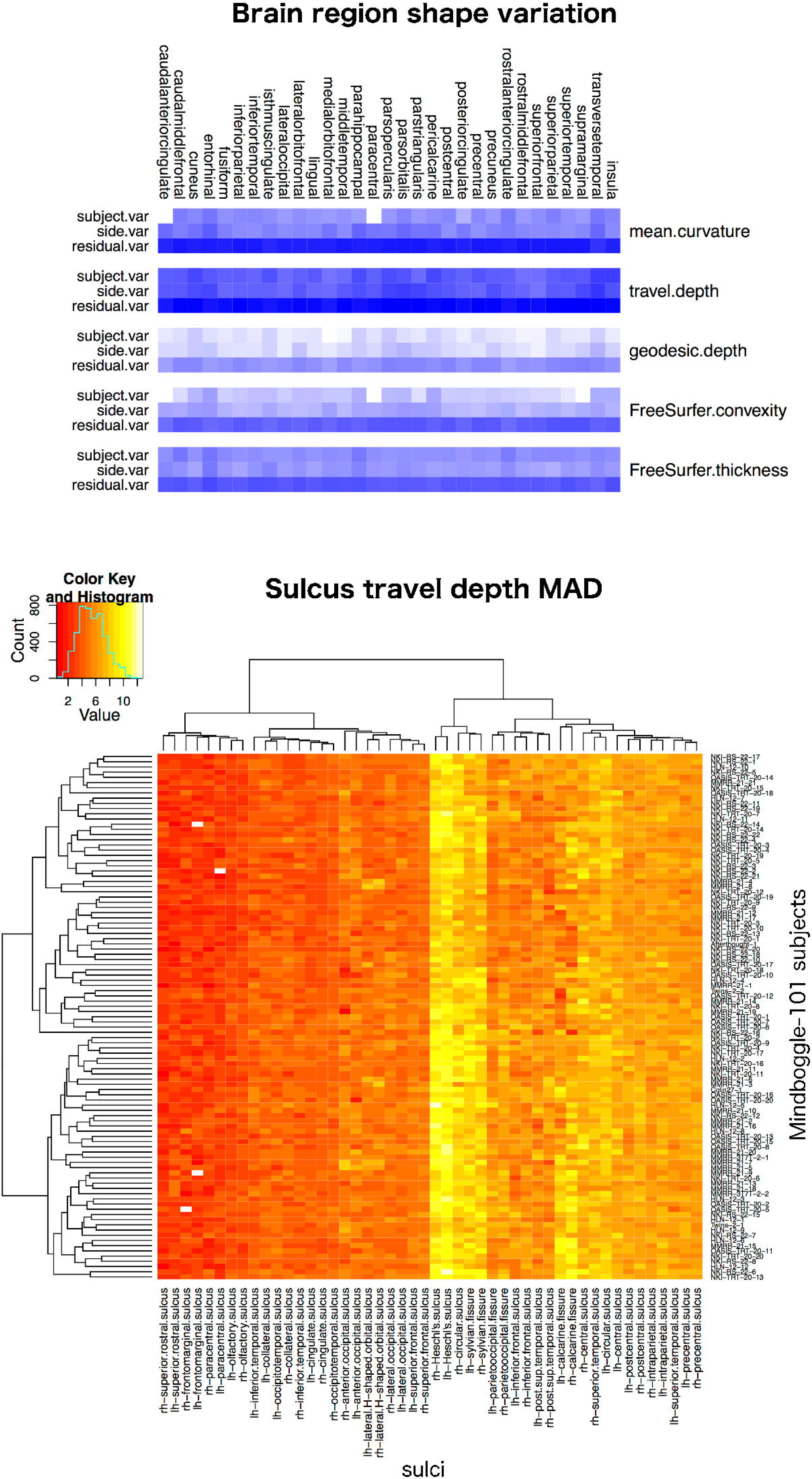
Brain shape variation in healthy humans. Top: Overview of the variance results for five shape measures computed on each of 31 manually labeled cortical regions (combined across both hemispheres for this figure) in the 101 Mindboggle-101 healthy human brains. The table shows the relative contributions of subject, hemisphere, and residual to describe the variability for each shape measure, with a greater contribution coded by a darker blue. For all shape measures and brain regions, most of the variability was concentrated in the residual. See **Supplement 2** for a description of the statistical models. Bottom: An example heatmap table containing 4,848 cells, where each cell is color-coded (increasing from red to yellow) to represent the median absolute deviation of travel depth values across all vertices in each of 48 sulcus surface meshes for the 101 subjects.

We repeated the same analysis as above on two scans acquired three years apart from hundreds of the ADNI participants (126 with Alzheimer’s, 199 healthy controls) as part of an international Alzheimer’s challenge (see Section 2.1: 2015 above) to see if we could find changes in brain shape measures that correlate with changes in ADNI-MEM cognitive scores over the course of three years. This resulted in the most detailed shape analysis of brains with Alzheimer’s disease ever conducted [118] (https://osf.io/d7hx8/). To identify shape measures associated with Alzheimer’s disease, we used the average of the ranks of the following tests in that study: Kolmogorov-Smirnov test to see if there was a difference between distributions at baseline and at three years, and correlation of change in shape and change in ADNI-MEM cognitive scores.

We found that healthy brains and brains with Alzheimer’s disease have similar shape statistical summaries, but changes in the following shape measures after a three-year interval were significantly correlated with changes in ADNI-MEM cognitive score:

- Volume for right caudal anterior cingulate and left: entorhinal, inferior parietal, (middle, superior) temporal, superior frontal, precuneus, and supramarginal gyri
- FreeSurfer thickness for left and right: entorhinal, fusiform, inferior parietal, (inferior, middle, superior) temporal, superior frontal, precuneus, and supramarginal gyri; left: (caudal middle/lateral, orbito/rostral middle) frontal, and pars triangularis gyri; right lingual gyrus
- Mean curvature for left and right rostral middle frontal gyri; left (middle, superior) temporal gyri; right inferior temporal gyri

## 4. Discussion

In this article, we have documented the Mindboggle open source brain morphometry platform and demonstrated its use in studies of shape variation in healthy and diseased humans. The number of different shape measures and the size of the populations make this the largest and most detailed shape analysis of human brains every conducted. There are many ways in which the open source software community can extend Mindboggle’s capabilities, and there are many possible applications for Mindboggle to brain and non-brain data. Here we will very briefly summarize this article’s results and point toward possible further evaluations and alternative approaches.

### 4.1 Summary of results

In this section we summarize the findings of our evaluations in the Results section. Shape measures are not independent of one another, and some related shape measures exaggerate values for certain morphological structures (such as geodesic vs. travel depth for the insula). Mindboggle’s `thickinthehead` cortical thickness measure is consistent across scans and across brains and generated values that are closer to published ranges of values than FreeSurfer or ANTs values. Mindboggle’s travel depth measure provides a more faithful representation of depth for shallow portions than FreeSurfer’s convexity measure. Mindboggle’s fundi are comparable to Gang Li’s fundi in terms of average proximity to manual label boundaries, but there was no clear winner in our evaluation of fundus extraction algorithms. Mindboggle’s shape measures are reasonably consistent across scans of the same brain, with some exceptions (such as entorhinal volume). We found that for the shape measures and populations we studied that shape differences between hemispheres were greater than shape differences between MRI scans of the same hemisphere, and that the variability within each brain hemisphere was higher than the variability between brain hemispheres in a participant or between participants. Finally, we reported which brain regions were significantly correlated with changes in ADNI-MEM cognitive score after a three-year interval as part of an international Alzheimer’s challenge.

### 4.2 Further evaluations and enhancements of Mindboggle

The Mindboggle software will continue to be subjected to evaluations of its algorithms as well as of its applicability to new datasets of healthy and diseased brains. Data exist to conduct evaluations of test/retest reliability and reproducibility [119, 120], with different imaging parameters [121], with genetic information [122], with heritability information [123], at higher field strengths [124], etc. Some data also exist for evaluating features such as sulcal pits [125]. Including different types of brain images can enable multivariate analyses, independent corroboration of morphology, and can even help to better interpret the factors that influence morphology [79]. We also intend to evaluate Mindboggle output by analyzing interactions among shape measures to find higher order morphological relationships with brain shape differences. There are many ways to enhance Mindboggle’s functionality and applicability to pathological brains. Taking advantage of different and multiple types of images, atlases, labels, features, and shape measures are clear ways to expand and improve Mindboggle, and the software was built using the Nipype framework specifically to enable modular and flexible inclusion of different algorithms. Even the inputs to Mindboggle can change to take advantage of promising new algorithms that combine surface reconstruction with whole-brain segmentation in a way that is more robust to white-matter abnormalities [126]. Use of probabilistic labels, features, and shape measures could lead to more careful interpretations of morphometry studies.

### 4.3 Alternative approaches to Mindboggle: Deep learning and beyond

The Mindboggle software extracts and identifies features for shape analysis. This approach is based on human-designed features (brain structure and label definitions and algorithmic implementations) and assumes the validity of the designed feature model. The tremendous success that machine learning (especially deep learning) approaches have had across domains [127] are strong evidence that such approaches may improve automated feature extraction, identification and labeling for features that a human would never consider designing. Machine learning has recently been demonstrated to recognize the multi-modal ‘fingerprint’ of cortical areas [24]. In 2011, we advocated a novel application of convolutional networks to build discriminative features and were able to demonstrate automated volumetric labeling of the cerebral cortex, without human intervention to build handcrafted features or to provide other prior knowledge [128]. At the time we had very limited training data (40 manually labeled brains), but with the Mindboggle-101 dataset, tables of shape statistics generated by Mindboggle, and with improved deep learning architectures, we may now be in a better position to apply deep learning to this problem. It may be helpful to explore ways in which priors and invariances can be modeled and integrated into deep learning approaches to reduce the amount of required training data and to integrate human expert information. This may be particularly beneficial for pathological conditions with tumors, lesions, and edemas, etc. that do not conform to a canonical reference brain or are difficult to obtain in sufficient quantities to train a deep learning algorithm. Indeed, thousands or millions of curated and labeled examples are usually required for deep learning algorithms, which points to the promise of unsupervised approaches that do not require expert feedback during training and can learn from messier data or from less data. Combining algorithmic approaches to feature extraction and morphometry with machine learning and unsupervised approaches has great potential applications in characterizing not just healthy human brain variation but in diagnosing, tracking, and predicting unhealthy conditions.

**Supporting Information**

**Document S1. Travel depth**

**Document S2. Variance components analysis of the shapes of 62 cortical regions in 101 human brains**

**Appendix S3. Mindboggle output directory tree**

## Acknowledgments

We sincerely thank everyone who has contributed to the Mindboggle open science project over the years, including Nolan Nichols, Oliver Hinds, Arthur Mikhno, Hal Canary, and Ben Cipollini. We also thank Gang Li, Denis Rivière, and Olivier Coulon for assistance with the fundus evaluation. Arno Klein would like to thank Deepanjana and Ellora for their continued patience and support, and dedicates this project to his mother and father, Karen and Arnold Klein.

Software and online resources that have benefitted the project include: GitHub.com, ReadtheDocs.org, and PyCharm for software development; Open Science Framework, Harvard Dataverse, and Synapse.org for public data storage; and D3, Bokeh, Paraview, and the Viridis colormap (https://github.com/bids/colormap) for visualization.

## Author Contributions

Arno Klein led the development of the project, organized the studies, was the main contributor to the software, and wrote the manuscript. Satrajit S. Ghosh contributed to the development of the software pipeline framework, has provided invaluable guidance over the years, and led the consulting group TankThink Lab’s involvement in the project. Forrest S. Bao wrote an early version of the software for extracting morphological features from brain surfaces and an open source Python version of Martin Reuter’s Laplace-Beltrami code. Joachim Giard wrote the open source C++ software for measuring depth and curvature along brain surfaces. Yrjö Häme explored hidden Markov measure field-based feature extraction. Eliezer Stavsky performed early explorations into surface feature-based label propagation. Noah Lee contributed to the original R01 proposal that supported Mindboggle’s research efforts, created an early graph-based database with interactive visualization, and implemented a deep learning approach to automate anatomical labeling. Martin Reuter helped to port his Laplace-Beltrami spectral shape measure to Python and evaluate the code. Elias Chaibub Neto performed statistical analyses of shape variation. Anisha Keshavan is helping to develop the ROYGBIV interactive online brain image viewer for visualizing Mindboggle output.

